# Adaptation delay causes a burst of mutations in bacteria responding to oxidative stress

**DOI:** 10.1101/2022.05.25.493234

**Authors:** Valentine Lagage, Victor Chen, Stephan Uphoff

## Abstract

Understanding the interplay between phenotypic plasticity and genetic adaptation is a long-standing focus of evolutionary biology. In bacteria, the oxidative stress response limits the formation of mutagenic reactive oxygen species (ROS) under diverse stress conditions. This suggests that the dynamics of the oxidative stress response are closely tied to the timing of the mutation supply that fuels genetic adaptation to stress. Here, we explored how mutation rates change in real-time in *Escherichia coli* cells during continuous hydrogen peroxide treatment in microfluidic channels. By visualising nascent DNA replication errors, we uncovered that sudden oxidative stress causes a burst of mutations. We developed a range of single-molecule and single-cell microscopy assays to determine how these mutation dynamics arise from phenotypic adaptation mechanisms. Signalling of peroxide stress by the transcription factor OxyR rapidly induces ROS scavenging enzymes. However, an adaptation delay leaves cells vulnerable to the mutagenic and toxic effects of hydroxyl radicals generated by the Fenton reaction. Resulting DNA damage is counteracted by a spike in DNA repair activities during the adaptation delay. Prior stress exposure or constitutive OxyR induction allowed cells to avoid the burst of mutations. Similar observations for alkylation stress show that mutation bursts are a general phenomenon associated with adaptation delays.

## Introduction

Phenotypic and genetic plasticity are at the root of the remarkable capacity of bacteria for adaptation. Stress responses induce phenotypic changes that protect bacteria against adverse conditions in the environment, thus opening a window of opportunity during which adaptive mutations may arise that provide long-term heritable stress resistance. The relative timing of phenotypic and genetic changes is critical to the process of adaptation. On the one hand, rapid phenotypic responses allow cells to survive transient stresses (“Bact. Stress Responses, Second Ed.,” 2011) without having to acquire permanent genetic changes which are rarely beneficial to the individual when the stress has passed (Robert et al., 2018). On the other hand, severe stress conditions can quickly drive a bacterial population to extinction. In such situations, protection from phenotypic responses is insufficient and population survival relies on evolutionary rescue via the rapid emergence of adaptive mutations (Bell, 2017). These mutations may be present as pre-existing genetic variation in a population due to spontaneous mutagenesis prior to the stress exposure or generated de-novo during stress. Understanding the interplay between phenotypic responses and genetic adaptation requires knowledge of how mutation rates change in real-time when cells become stressed and how these dynamics relate to the underlying molecular mechanisms of adaptation. However, experimental evidence for the actual temporal order of phenotypic and genetic changes during stress adaptation is lacking because conventional methods for detecting mutations do not resolve phenotypic responses, and vice versa. This has now become possible owing to a new live-cell microscopy approach to measure DNA replication error rates while simultaneously monitoring cell growth, morphology, and gene expression dynamics (Robert et al., 2018; Uphoff, 2018).

Amongst the most common and harmful stress factors that threaten bacteria are reactive oxygen species (ROS), which are constantly formed inside cells as a byproduct of their metabolism (Imlay, 2008). ROS levels increase under various environmental conditions. For example, sudden bursts of ROS are generated by immune cells to fight bacterial infections (Fang et al., 2016), when bacteria are exposed to bactericidal antibiotics (Giroux et al., 2017; Hong et al., 2019; Kohanski, Dwyer, et al., 2010) or radiation (Imlay, 2015), and during competition with other bacterial species (Dong et al., 2015). Hydrogen peroxide (H_2_O_2_) is one of the major ROS and can cause damage to proteins, lipids, and DNA through the formation of hydroxyl radicals (HO) via the Fenton reaction with intracellular iron (Imlay, 2008). Purine nucleobases are particularly prone to oxidation, resulting in pre-mutagenic lesions which can form mismatched base pairs during DNA replication, ultimately leading to mutations if not repaired (Bjelland & Seeberg, 2003). To counter this omnipotent threat to genome stability, dedicated enzymes of the DNA Base Excision Repair (BER) pathway revert pre-mutagenic oxidative lesions before they turn into mutations (Friedberg et al., 2005). As a primary defence against ROS toxicity, many bacteria rely on catalase and peroxiredoxin enzymes to scavenge intracellular H_2_O_2_ (Imlay, 2008). Although the specific mechanisms vary between bacterial species, the expression of ROS scavenging enzymes is generally regulated via redox-sensitive transcription factors such as OxyR, which control oxidative stress responses that substantially increase ROS tolerance (Storz et al., 1990).

Over time, prolonged oxidative stress leads to the selection of adaptive mutations in a variety of targets that affect iron homeostasis and cell motility (Rodríguez-Rojas et al., 2020), increase the expression of catalase (Xiaojing Li et al., 2014) or lead to constitutive OxyR induction (Anand et al., 2020; J & J, 2014). The mutagenic effects of ROS are thought to accelerate the evolution of antibiotic resistance and host adaptation (Kohanski, DePristo, et al., 2010; Long et al., 2016; H. Wang et al., 2018; Weitzman & Stossel, 1981). Interestingly, the mutation supply driving such genetic adaptation was shown to depend on the temporal pattern of ROS exposure (Rodríguez-Rojas et al., 2020), suggesting that mutation rates may be dynamically modulated by changes in the stress level and the cellular stress responses.

## Results

### Monitoring adaptation of *E. coli* to H_2_O_2_ treatment in microfluidic chips

We set out to monitor the timing of phenotypic adaptation and mutagenesis of *E. coli* cells exposed to H_2_O_2_ via live-cell microscopy. Conventional bulk culture assays typically employ a single H_2_O_2_ bolus which decays in the medium over time due to efficient ROS scavenging by the bacteria. Because of this, induction of prolonged oxidative stress had typically required treatment with very high (~ millimolar) H_2_O_2_ concentrations, which are outside the range experienced by bacteria in nature (Xin Li & Imlay, 2018). To avoid this, and to uncouple the dynamics of the bacterial responses from changes in the H_2_O_2_ concentrations in the media, we imaged cells growing inside the “mothermachine” microfluidic device (Uphoff, 2018; P. Wang et al., 2010). Individual cells are maintained within µm-sized channels under exponential growth conditions over tens of generations and supplied with a continuous inflow of fresh media and H_2_O_2_ treatment at a constant concentration (Fig. 1A). The individual cells at the closed end of each growth channel (named “mother cells”) can be monitored continuously, while their daughter cells are pushed towards the open ends of the channels and washed out with the media outflow. We used time-lapse epifluorescence microscopy to measure cell elongation rates (change in cell length) and generation times (interval between consecutive cell divisions) via automated segmentation of the cell shape aided by a cytoplasmic mKate2 fluorescent protein. An experiment typically monitored 500-1000 individual mother cells, each in a separate growth channel. After a period of unperturbed growth for several generations, we switched the media inflow to expose cells to a constant concentration of H_2_O_2_ that is continuously replenished by the fluidics system. H_2_O_2_ crosses the *E. coli* cell envelope readily (Seaver & Imlay, 2001), and the elongation rate dropped two-fold immediately after the start of 100 µM H_2_O_2_ treatment (Fig. 1B). We observed a partial inhibition of cell elongation for H_2_O_2_ concentrations as low as 25 µM, demonstrating that physiologically-relevant H_2_O_2_ concentrations are sufficient to cause detrimental oxidative stress (Fig. S1). At 500 μM H_2_O_2_ and above, cell elongation ceased almost completely, whereas the growth inhibition was only transient at milder treatments and elongation rates recovered to pre-treatment rates after an adaptation lag period. The decreased rate of cell elongation led to a delay in the cell cycle during the adaptation lag (Fig. 1D, Fig. S2). The adaptation lag was also marked by a transient period of increased cell mortality, which manifested either as irreversible cessation of elongation, sudden cell lysis, or extreme filamentation that caused the escape of the cell from the growth channel (Fig. 1B and Fig. S3). Due to the constant treatment, the recovery of cell elongation and viability cannot be attributed to a decay of H_2_O_2_ in the medium, but rather reflect the phenotypic adaptation that enables unperturbed growth in the presence of H_2_O_2_. However, an initial lag period before adaptation leaves the bacteria vulnerable to the toxic effects of H_2_O_2_ which have been attributed to the oxidation of DNA and proteins (Ezraty et al., 2017) and a shutdown of protein synthesis (Zhong et al., 2015; Zhu & Dai, 2019).

**Fig. 1:**
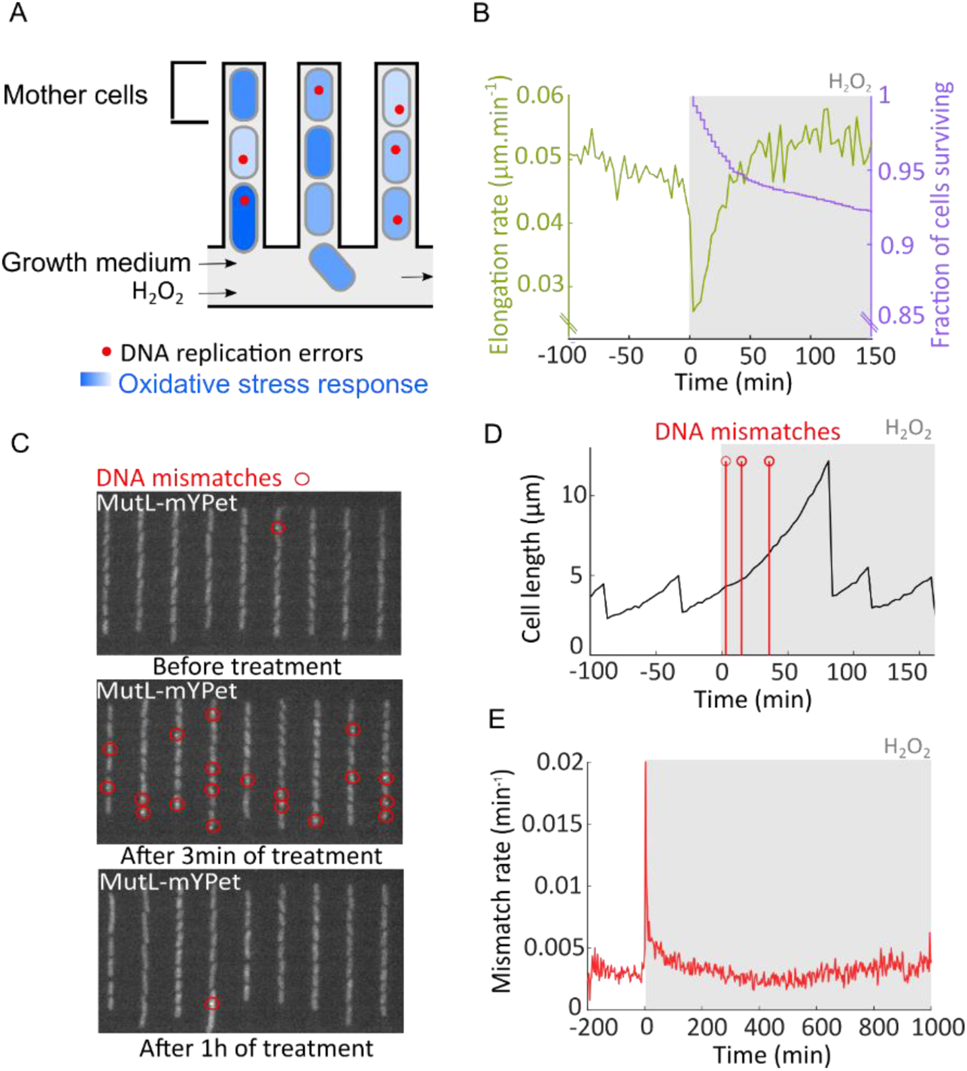
H_2_O_2_ treatment causes a burst of mutation. (A) Single-cell microscopy monitors DNA replication errors and fluorescent gene expression reporters for the oxidative stress response during constant H_2_O_2_ treatment using microfluidics. The “mother cells” are located at the closed end of each growth channel and can be observed continuously over multiple generations. (B) Cell elongation rate (green) and fraction of cells surviving (purple) in the microfluidic chip during constant treatment with 100 µM H_2_O_2_ added at time 0 (6657 cells, 8 experiments). (C) Snapshots of MutL-mYPet foci (red circles) marking DNA mismatches in cells before treatment, and after 3 min and 1 hour of constant treatment with 100 µM H_2_O_2_. (D) Example time trace of one mother cell growing and dividing before and during constant treatment with 100 µM H_2_O_2_ added at time 0. Red markers indicate DNA mismatch events. (E) Rate of DNA mismatches per cell per minute before and during constant treatment with 100 µM H_2_O_2_ (6655 cells, 8 experiments).

### Sudden oxidative stress causes a burst of mutations

We next asked how the timing of phenotypic adaptation relates to the rates of genetic change during oxidative stress. To this end, we employed an imaging-based method to monitor mutagenesis in live bacteria (Robert et al., 2018; Uphoff, 2018). Point mutations arise during DNA replication due to the erroneous incorporation of nucleotides that do not match the DNA template sequence. Certain chemical modifications of bases on the template strands are highly mutagenic due to their propensity to form DNA mismatches. While most DNA replication errors are corrected by the DNA mismatch repair (MMR) system, a small fraction of mismatches (1%) escapes repair and is converted into stable point mutations during the next replication cycle (Long et al., 2018). Fusion of the MMR enzyme MutL to the fluorescent protein mYPet yields a visual reporter for DNA replication errors in live cells, marked by the assembly of MutL-mYPet molecules in foci at a DNA mismatch (Elez et al., 2010; Uphoff, 2018). Notably, the frequency of MutL-mYPet foci reports on the rate of DNA replication errors, irrespective of whether a DNA mismatch ultimately turns into a mutation or is repaired. Nevertheless, it was shown that the rate of MutL-mYPet foci (mismatch rate) in a cell population correlates with the frequency of genomic mutations, validating its use as a mutation reporter (Vincent & Uphoff, 2021). Because DNA mismatches are rare and transient, we monitored thousands of cells in order to obtain a reliable estimate of the instantaneous mismatch rate per cell per minute. As seen before (Robert et al., 2018; Uphoff, 2018), the mismatch rate per cell per minute was stable over time in untreated cells at on average 0.0029 mismatches cell^−1^ min^−1^ (Fig. 6J and Fig. S4).

Treatment with 100 μM H_2_O_2_ caused a sudden ~10-fold increase in the mismatch rate per cell per minute (Fig. 1C). Single cells often showed multiple mismatch events shortly after the start of treatment (Fig. 1D). The mismatch rate per cell per minute averaged over a total of 6655 cells revealed a sharp mutagenesis burst with a duration of ~12 min and a maximum rate of 0.021 mismatches cell^−1^min^−1^ at 4.5 min after treatment started (Fig. 1E). The mutagenesis burst also occurred in the daughter cells located above the mother cell in the growth trenches (Fig. S5). No MutL-mYPet foci were detected in cells with a *mutS* gene deletion either before or after H_2_O_2_ exposure (Fig. S6). As MutS is required for recognition of DNA mismatches and the recruitment of MutL, this control confirms that MutL-mYPet foci genuinely show DNA mismatches. The maximum of the mutagenesis burst saturated with increasing H_2_O_2_ concentration (Fig. S1B), indicating that the formation of mutagenic lesions or their conversion into DNA mismatches become limited when cell growth is strongly inhibited by H_2_O_2_. Our observation that the mutagenesis burst also occurs at H_2_O_2_ concentrations that inhibit growth completely (>200 µM in microfluidic culture, Fig. S1B) indicates that it may contribute to the acquisition of mutations that enable evolutionary rescue during lethal stress (Rodríguez-Rojas et al., 2020).

Reactive oxygen species cause a variety of DNA lesions (Bjelland & Seeberg, 2003). The MMR pathway does not efficiently revert all types of DNA replication errors, such as G-A, C-C, and G-C mispairs (Brown et al., 2001), and it is hence expected that these mismatches cannot be detected using the MutL-mYPet reporter (Robert et al., 2018; Uphoff, 2018). To confirm the existence of the mutations burst using a different method, we measured the frequency of rifampicin-resistant colonies which is commonly used as a readout for genomic mutation rates (Fowler et al., 2003). We treated bulk cultures with 1 mM H_2_O_2_ for different length of time. The frequency of rifampicin-resistant colonies increased significantly for 5 min of treatment but there was no further increase when cultures were treated for 12 min or longer. Hence, all mutations occured in the initial 5 min of treatment. Although these measurements lack the temporal resolution of the MutL-mYPet reporter, the results of both methods are consistent and demonstrate that H_2_O_2_-induced mutagenesis is confined to a brief interval after the onset of treatment (Fig. S7).

### The burst of mutations coincides with a delay in the oxidative stress response

*E. coli* adapts to H_2_O_2_ by induction of catalase, peroxidase, glutaredoxin 1, and thioredoxin radical scavenging enzymes, which are under the control of the transcription factor OxyR (Storz et al., 1990). As expected, cells with a Δ*oxyR* gene deletion were hypersensitive to H_2_O_2_ and failed to adapt to the treatment over time (Fig. 2A-B). In contrast to the sharp mutagenesis burst seen in the wild-type (WT), the lack of adaptation in the Δ*oxyR* strain led to a gradual and sustained increase of the mismatch rate per cell per minute in the presence of H_2_O_2_, although the observation time was limited by rapid death of Δ*oxyR* cells (Fig. 2C). Furthermore, a Δ*katG* mutant lacking the OxyR-inducible catalase showed complete loss of growth with 100 μM H_2_O_2_ and an elevated mutagenesis burst compared to the WT (Fig. 2D-F). We saw no difference in the basal mismatch rate per cell per minute for Δ*katG* compared to the WT (Fig. 2F), consistent with the alkyl hydroperoxidase AhpCF being the main scavenging enzyme of endogenous H_2_O_2_ in untreated cells (Costa & Imlay, 2001). To address how the observed timing of phenotypic adaptation and mutagenesis relate to the gene expression dynamics of the OxyR response, we imaged transcriptional reporters for the catalase promoter PkatG fused to the fluorescent protein SCFP3A (Balleza et al., 2018) and the glutaredoxin 1 promoter PgrxA-SCFP3A, expressed from low-copy number plasmids (Zaslaver et al., 2006). As expected, both reporters were strongly induced during H_2_O_2_ treatment (Fig. 2G-H, Fig. S1C), and no gene induction occurred in the Δ*oxyR* strain (Fig. S8). Both reporters showed an induction peak shortly after treatment before dropping to a constant intermediate expression level. This can be explained by a high intracellular H_2_O_2_ concentration causing a strong upregulation of scavenging enzymes in naïve cells. As the scavenging capacity increases, the resulting drop in H_2_O_2_ inside cells leads to a steady-state expression where the stress and OxyR response are balanced. The fluorescence signal of PgrxA and PkatG reporters reached half-maximal intensity after 15 (median value) min and 12 min (median value) of treatment, respectively (Fig, 2H, Fig. S9). This short lag time of the stress response aligns precisely with the duration of the mutagenesis burst. Indeed, simultaneous imaging of the gene expression reporters and MutL-mYPet foci in the same cells revealed that the mutagenesis burst starts to decrease as soon as the OxyR reporter signals increase (Fig. 2I). Notably, cells that acquire adaptive mutations early during the onset of stress would still rely on stress responses to bridge the phenotypic delay before the adapted phenotype is expressed (Sun et al., 2018).

**Fig. 2:**
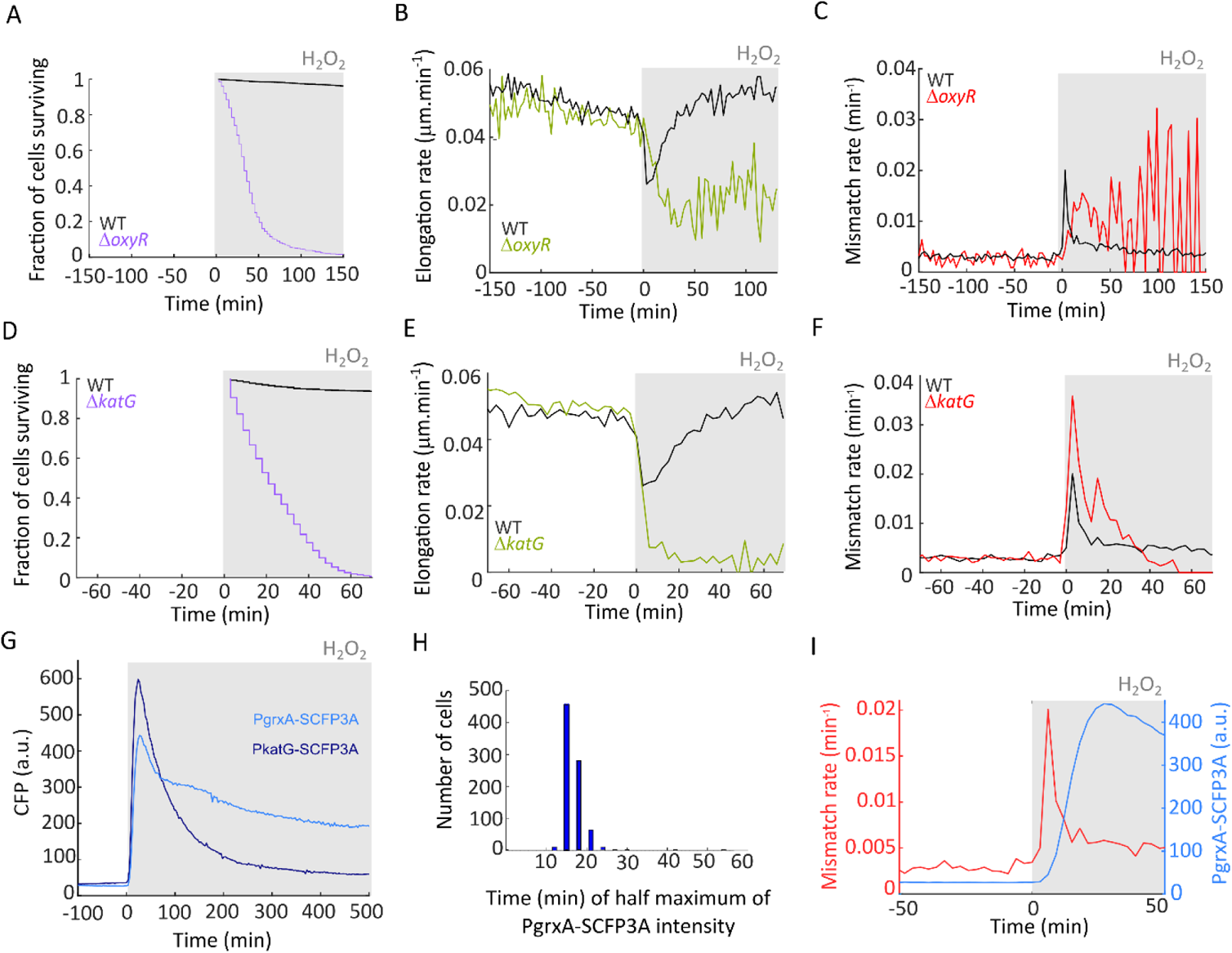
The OxyR response and induction of scavenging enzymes are required for adaptation to constant H_2_O_2_ treatment. (A-C) OxyR mutant (Δ*oxyR*). (A) Fraction of cells surviving after start of treatment (734 cells, 3 experiments) (B) Cell elongation rate (668 cells, 3 experiments). (C) Rate of DNA mismatches per cell per minute (728 cells, 3 experiments). (D-F) Catalase deletion mutant (Δ*katG*) before and during constant treatment with 100 µM H_2_O_2_. (D) Fraction of cells surviving after start of treatment (3040 cells, 3 experiments). (E) Cell elongation rate (2787 cells, 3 experiments). (F) Rate of DNA mismatches per cell per minute (3137 cells, 3 experiments). (G) PgrxA-SCFP3A (1848 cells, 2 experiments) and PkatG-SCFP3A (988 cells, 1 experiment) fluorescence intensity before and during constant treatment with 100 µM H_2_O_2_ of WT cells. (H) Histogram of the lag time distribution for PgrxA-SCFP3A to reach half-maximal intensity after start of 100 µM H_2_O_2_ treatment (878 cells, 1 experiment). (I) Joined plot of the DNA mismatch rate per cell per minute (Fig. 1E) and PgrxA-SCFP3A fluorescence intensity (Fig. 2G).

### Single-molecule tracking shows the dynamics of OxyR activation inside cells

Our observation that an adaptation delay causes a burst of mutagenesis during H_2_O_2_ exposure led us to examine the timing of H_2_O_2_ sensing and signalling upstream of the gene expression response. For this, we developed a novel approach based on single-molecule tracking of OxyR (Fig. 3A-B). OxyR senses H_2_O_2_ via reversible oxidation that leads to the formation of a disulphide bond which increases its binding affinity for target gene promoters to activate their transcription (Lee et al., 2004; Zheng et al., 1998). We reasoned that promoter binding should be detectable by measuring the mobility of OxyR in cells. To this end, we created a stable HaloTag fusion of OxyR for single-molecule fluorescence imaging (Fig. S10). The *E. coli* strain expressing OxyR-Halo from the native endogenous locus showed the same tolerance to H_2_O_2_ as WT cells, in contrast to the hypersensitivity of a Δ*oxyR* mutant (Fig. S11). Furthermore, induction of the PgrxA-SCFP3A reporter with 100 μM H_2_O_2_ was unaffected by the HaloTag fusion to OxyR (Fig. S12). These tests demonstrate that the OxyR-Halo fusion maintains full functionality. We performed single-molecule tracking of OxyR-Halo labelled with the cell-permeable dye TMR (Banaz et al., 2019). In untreated cells, most OxyR molecules exhibited random diffusive motion, whereas H_2_O_2_ treatment led to the appearance of a distinct population of immobile molecules (Fig. 3B-C). By measuring the diffusion coefficient (D), we quantified the fraction of immobile molecules relative to the diffusing pool of molecules as a readout for H_2_O_2_ sensing by OxyR. The immobile OxyR population peaked at 5 min after the start of H_2_O_2_ treatment, which was the earliest time point we were able to record with this method. Oxidised OxyR is deactivated via reduction by glutaredoxin 1 (Zheng et al., 1998), leading to a decay in the population of promoter-bound OxyR when the scavenging system has lowered the intracellular H_2_O_2_ concentration (Fig. 3C). The half-life of the immobile OxyR population was ~15 min and OxyR binding returned to the basal level after ~30 min of constant H_2_O_2_ treatment, consistent with previous bulk measurements (Tao, 1999). To relate the dynamics of OxyR binding to the transcription of its target genes, we measured the PgrxA and PkatG promoter activities by computing the time-derivatives of the SCFP3A reporter signals (Fig. 3D-E, Fig. S13A). The period of increased OxyR binding coincided with a spike in the promoter activities (Fig. 3F, Fig. S13B). Together, these direct measurements show that maximal activation of OxyR occurs within a few minutes of constant H_2_O_2_ exposure and causes a rapid increase in the transcription rate of oxidative stress response genes.

**Fig. 3:**
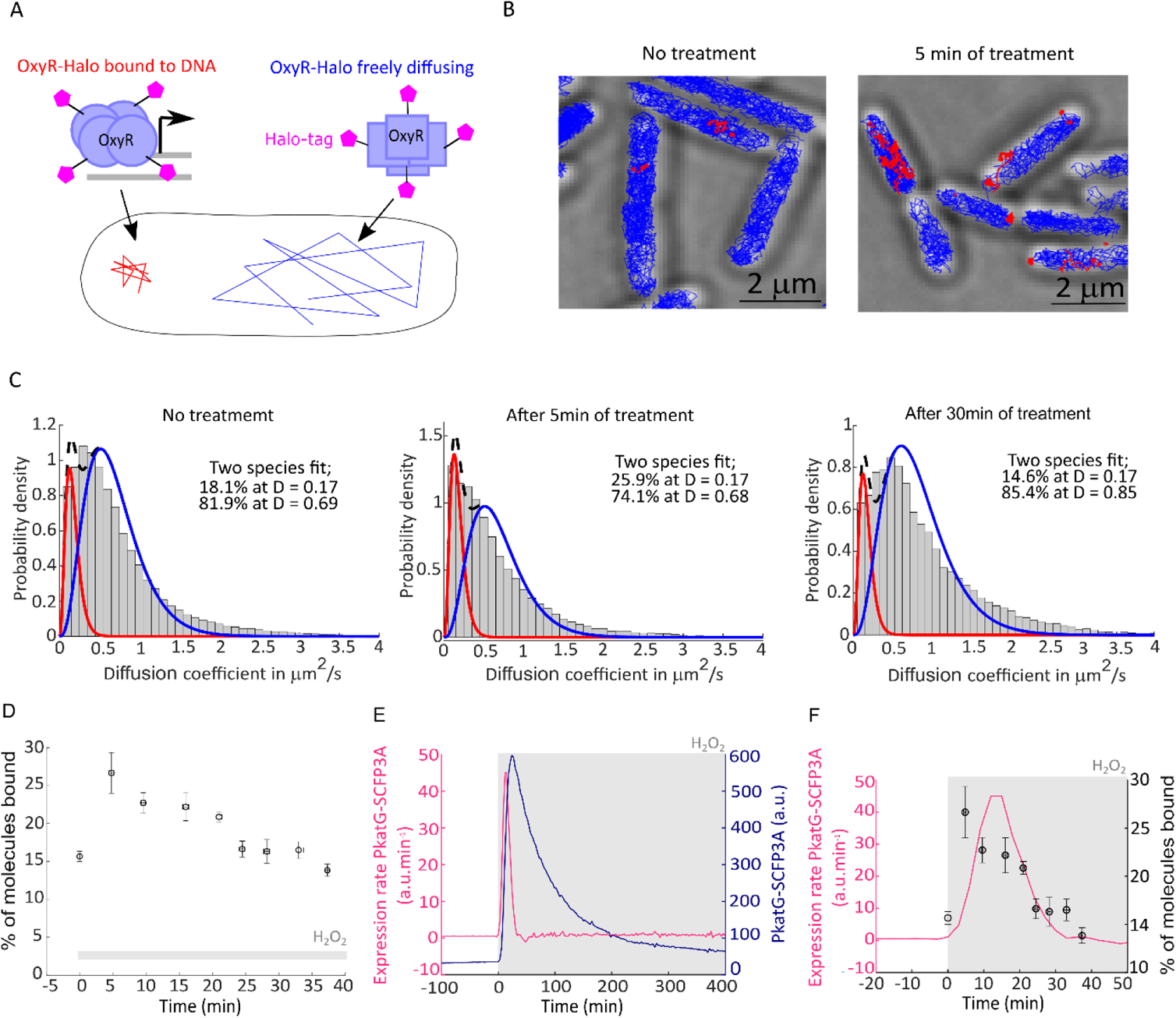
OxyR induces target genes within <5 minutes after the start of H_2_O_2_ treatment. (A) Schematic of single-molecule tracking to detect DNA-binding of OxyR-Halo based on the diffusion coefficient. (B) Representative tracks of bound (red, diffusion coefficient (D) < 0.17 µm^2^/s) and diffusing OxyR-Halo molecules (blue, D > 0.17 µm^2^/s) in untreated cells (left) and after 5 min of 100 µM H_2_O_2_ treatment (right). Scale bar 2 µm. (C) Distribution of diffusion coefficients of OxyR-Halo in untreated cells (Left, 83271 tracks, 3 experiments), after 5 min of treatment (Middle, 27213 tracks, 3 experiments), and after 30 min of treatment (Right, 16186 tracks, 3 experiments) with 100 µM H_2_O_2_. Distributions were fitted to quantify the relative abundances of bound molecules (with D = 0.17 µm^2^/s) and diffusing molecules (with D ~ 0.69 – 0.85 µm^2^/s).(D) Percentage of bound OxyR-Halo molecules at different time points after 100 µM H_2_O_2_ treatment (mean ± SEM, 3 experiments). (E) Expression rate of PkatG-SCFP3A (pink) and PkatG-SCFP3A intensity (dark blue) with 100 µM H_2_O_2_ treatment (988 cells, 1 experiment). (F) Expression rate of PkatG-SCFP3A (from panel E, pink) and percentage of bound OxyR-Halo molecules (from panel D, black).

### Constitutive OxyR response and priming adaptation prevent the mutagenesis burst

To test whether the transient delay of the OxyR response is indeed responsible for the sharp mutagenesis burst, we performed experiments under conditions in which the delay was abolished. First, we generated a strain with a constitutive OxyR response by elevating endogenous H_2_O_2_ levels with a deletion of the alkyl hydroperoxidase gene *ahpC* (Seaver & Imlay, 2001). This strain showed increased basal expression of the PgrxA reporter before treatment (Fig. 4A) and its elongation rate and survival were less affected after addition of 100 μM H_2_O_2_ treatment than WT cells (Fig, 4B-C, Fig. S14). Furthermore, the mutagenesis burst was completely absent (Fig. 4D). Instead, the mismatch rate per cell per minute increased gradually during constant H_2_O_2_ treatment in Δ*ahpC* cells, indicating that the alkyl hydroperoxidase contributes also to scavenging of exogenous H_2_O_2_. Second, exposing cells to sub-lethal concentrations of H_2_O_2_ is known to increase their survival of subsequent treatments (Imlay, 2008; Rodríguez-Rojas et al., 2020). To test if priming adaptation can also protect against the mutagenesis burst, we pre-treated cells with a low dose of H_2_O_2_ (25 μM) for 4 hours before exposure to the higher dose (100 μM). The pre-treatment induced expression of the PgrxA reporter to an intermediate level (Fig. 4E). Primed cells showed no reduction in elongation rate and no drop in survival when exposed to 100 μM H_2_O_2_ (Fig. 4F-G). Moreover, the mutagenesis burst was abolished in primed cells (Fig. 4H). Therefore, the lack of an adaptation delay due to a step-wise increase in oxidative stress level is sufficient to avoid the mutagenesis burst. This is in agreement with a reduction in mutation frequency as seen by genome sequencing of primed cells treated with H_2_O_2_ (Rodríguez-Rojas et al., 2020).

**Fig. 4:**
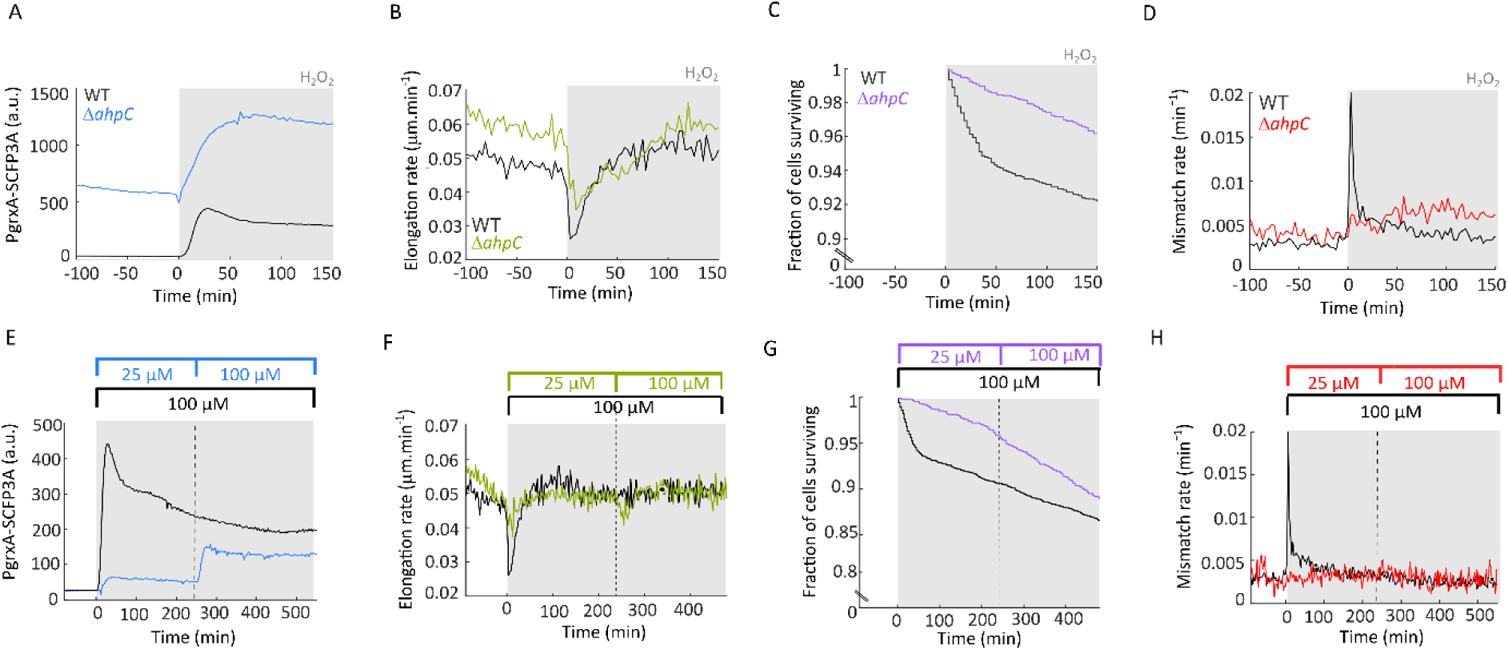
Constitutive OxyR response and priming-adaptation prevent the mutagenic and toxic effects of sudden H_2_O_2_ exposure. (A-D) Oxidative stress response with 100 µM H_2_O_2_ treatment of Δ*ahpC* deletion mutant (coloured lines) compared to WT (black lines). (A) PgrxA-SCFP3A intensity (1433 cells, 2 experiments). (B) Cell elongation rate (3015 cells, 5 experiments). (C) Fraction of cells surviving after start of treatment (2795 cells, 4 experiments). (D) Rate of DNA mismatches per cell per minute (3360 cells, 5 experiments). (E-H) Oxidative stress response of WT cells pre-treated with 25 µM H_2_O_2_ for 4 hours followed by constant treatment with 100 µM H_2_O_2_. Response to 100 µM H_2_O_2_ without pre-treatment is shown for comparison (black). (E) PgrxA-SCFP3A intensity (605 cells, 1 experiment). (F) Cell elongation rate (1830 cells, 3 experiments). (G) Fraction of cells surviving after start of treatment (1377 cells, 2 experiments). (H) Rate of DNA mismatches per cell per minute (2020 cells, 3 experiments).

### DNA repair dynamics during oxidative stress

Our results indicate that the low rates of replication errors after an initial mutagenesis burst can be attributed to a reduction in intracellular H_2_O_2_ concentration following the upregulation of ROS scavenging enzymes. However, it is also plausible that DNA damage levels remain elevated even after adaptation but that DNA repair mechanisms efficiently remove mutagenic lesions before they lead to replication errors. If so, we would expect that DNA damage levels and DNA repair activities are sustained after adaptation. To test this, we measured the expression dynamics of the SOS response which is induced by oxidative DNA damage that stalls DNA replication and generates DNA breaks (Imlay, 2008; Kreuzer, 2013). H_2_O_2_ treatment induced the expression of a PrecA-SCFP3A reporter for the SOS response (Fig. 5A). However, after an initial pulse of expression at 100 min, the reporter signal returned to the basal level after ~4-5 hours of constant treatment. Therefore, DNA damage levels do not remain elevated after adaptation.

**Fig. 5:**
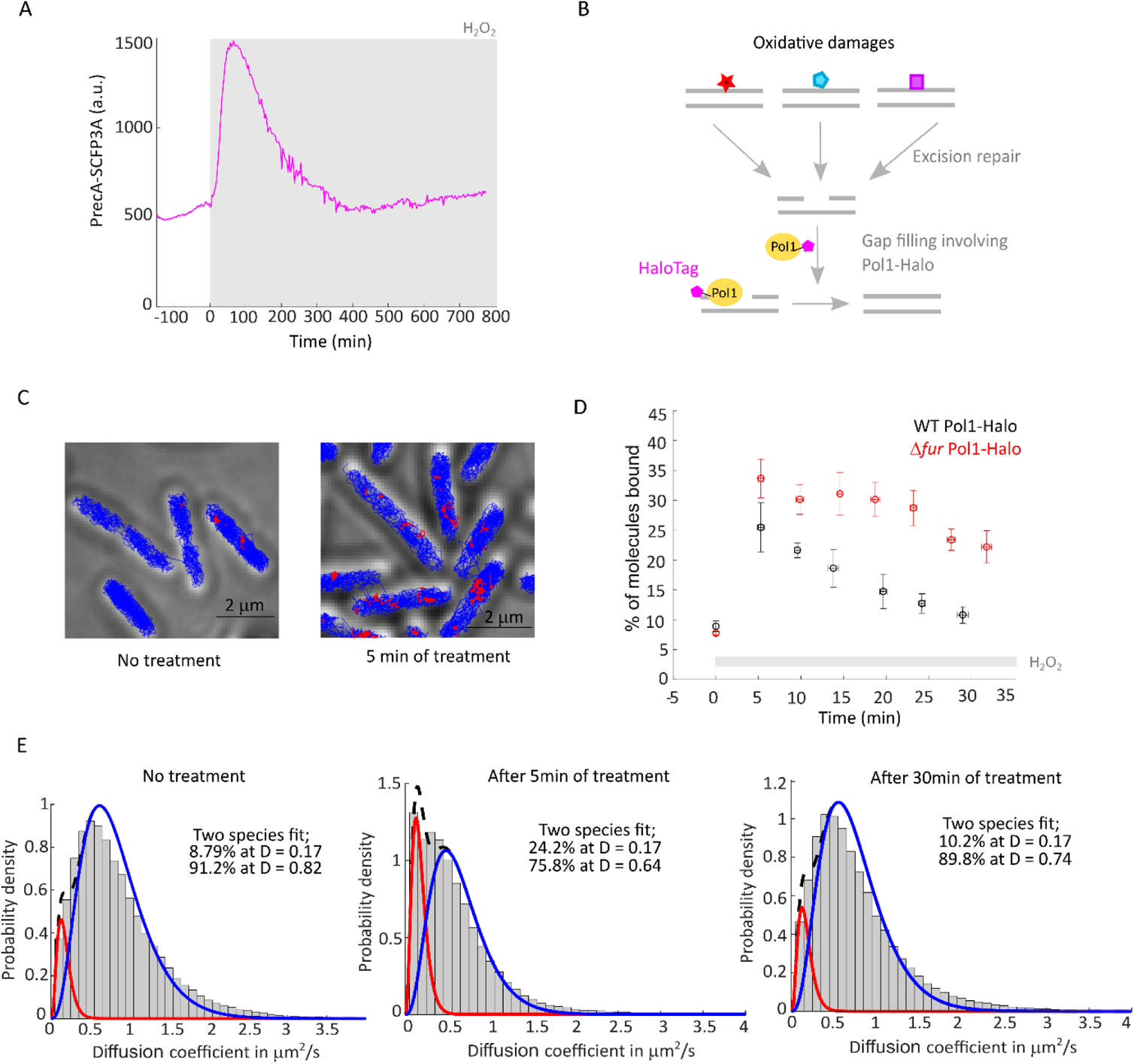
Dynamics of DNA damage response and DNA repair during oxidative stress adaptation. (A) SOS response reporter PrecA-SCFP3A expression with 100 µM H_2_O_2_ treatment (861 cells, 1 experiment). (B) Single-molecule tracking of Pol1-Halo as a reporter for repair of oxidative DNA lesions. Immobile molecules reflect DNA-bound Pol1. (C) Representative tracks of bound (red, D < 0.17 µm^2^/s) and diffusing Pol1-Halo molecules (blue, D > 0.17 µm^2^/s) in untreated cells (left) and after 5 min of 100 µM H_2_O_2_ treatment (right). Scale bar 2 µm. (D) Percentage of bound Pol1-Halo molecules in non-treated cells (time 0) and in response to 100 µM H_2_O_2_ treatment for WT cells (mean ± SEM, 3 experiments) and Δ*fur* mutant cells (mean ± SEM, 4 experiments). (E) Distribution of diffusion coefficients of Pol1-Halo without treatment (83613 tracks, 3 experiments), and after 5 min (98044 tracks, 3 experiments) and 30 min (112780 tracks, 3 experiments) of 100 µM H_2_O_2_ treatment. Distributions were fitted to quantify the relative abundances of bound molecules (with D = 0.17 µm^2^/s) and diffusing molecules (with D ~ 0.64 – 0.82 µm^2^/s).

Different types of oxidative DNA lesions are targeted by a variety of DNA repair enzymes via the BER and Nucleotide Excision Repair (NER) pathways. To directly measure the timing of overall repair of oxidative lesions during adaptation to H_2_O_2_, we chose DNA polymerase I (Pol1) as a universal reporter because it acts on DNA gaps which are a common intermediate in both BER and NER (Friedberg et al., 2005). Single-molecule tracking of a functional Pol1-Halo fusion (Banaz et al., 2019; Uphoff et al., 2013) (Fig. 5B) showed that the population of immobile molecules increased after H_2_O_2_ treatment, with the highest binding activity seen at the earliest recorded time point (5 min) (Fig. 5C-E). Pol1 did not show sustained binding activity during constant H_2_O_2_ treatment (Fig. 5C-E). Together, these results demonstrate that the induction of the oxidative stress response effectively protects cells against H_2_O_2_ present in their environment, reducing the formation of oxidative DNA damage down to the basal level within 30 min of continuous oxidative stress. This explains observations from genetic studies that deletion of BER and NER enzymes does not sensitise cells to killing by H_2_O_2_ (Hoff et al., 2021).

### Perturbation of intracellular iron levels modulates basal mutagenesis and the mutagenesis burst induced by H_2_O_2_

The toxic and mutagenic effects of H_2_O_2_ are attributed to the Fenton reaction, which couples the oxidation of ferrous iron to the generation of highly reactive hydroxyl radicals (Imlay, 2008). Consequently, regulation of iron homeostasis is a central feature of oxidative stress responses (Troxell & Hassan, 2013). To test the importance of iron for the observed mutagenesis burst, we added the iron chelator 2,2’-dipyridyl (DP) to the growth media before and during H_2_O_2_ treatment. We used a moderate concentration of DP to reduce free iron concentrations without perturbing cell growth rates (Fig. S15, Fig. 6A). The magnitude of the mutagenesis burst and the effect of H_2_O_2_ on cell mortality were both significantly reduced by DP (Fig. 6B-C, Fig. 6J-L). We still observed a transient reduction in cell elongation after H_2_O_2_ addition, but DP accelerated the adaptation and return to normal growth (Fig. 6A). Next, to decrease intracellular iron levels, we deleted the *tonB* gene, which is required for iron import (Chakraborty et al., 2007). This mutant had the same generation time (Fig. S15) and elongation rate as the WT but exhibited a significantly reduced rate of DNA mismatches per cell per minute even before treatment (Fig. 6D-F, Fig. 6J), showing that intracellular iron contributes substantially to spontaneous mutation in optimal growth conditions. Furthermore, the mutagenesis burst was completely absent in Δ*tonB* cells treated with H_2_O_2_, and we saw no reduction of growth or survival after treatment (Fig. 6D-F, Fig. 6K-L). However, the OxyR response was induced to the same level in WT, Δ*tonB*, and DP-treated cells, confirming that inhibition of the Fenton reaction does not affect H_2_O_2_ levels and oxidative stress response signalling (Fig. S16). To probe whether mutagenesis and toxic effects were limited by the amount of intracellular iron, we deleted the *fur* gene, which leads to increased iron import (Touati et al., 1995) Cell elongation and survival were more strongly attenuated during the adaptation delay for Δ*fur* cells (Fig. 6G-H). Although the height of the mutagenesis burst was unaffected by *fur* deletion, mismatch rates per cell per minutes remained elevated post adaptation and the OxyR response was induced at higher level than in WT cells (Fig. 6I, Fig. 6K-L, Fig. S16). Furthermore, the DNA repair activity of Pol1-Halo was increased and prolonged in the Δ*fur* mutant (Fig. 5D).

**Fig. 6:**
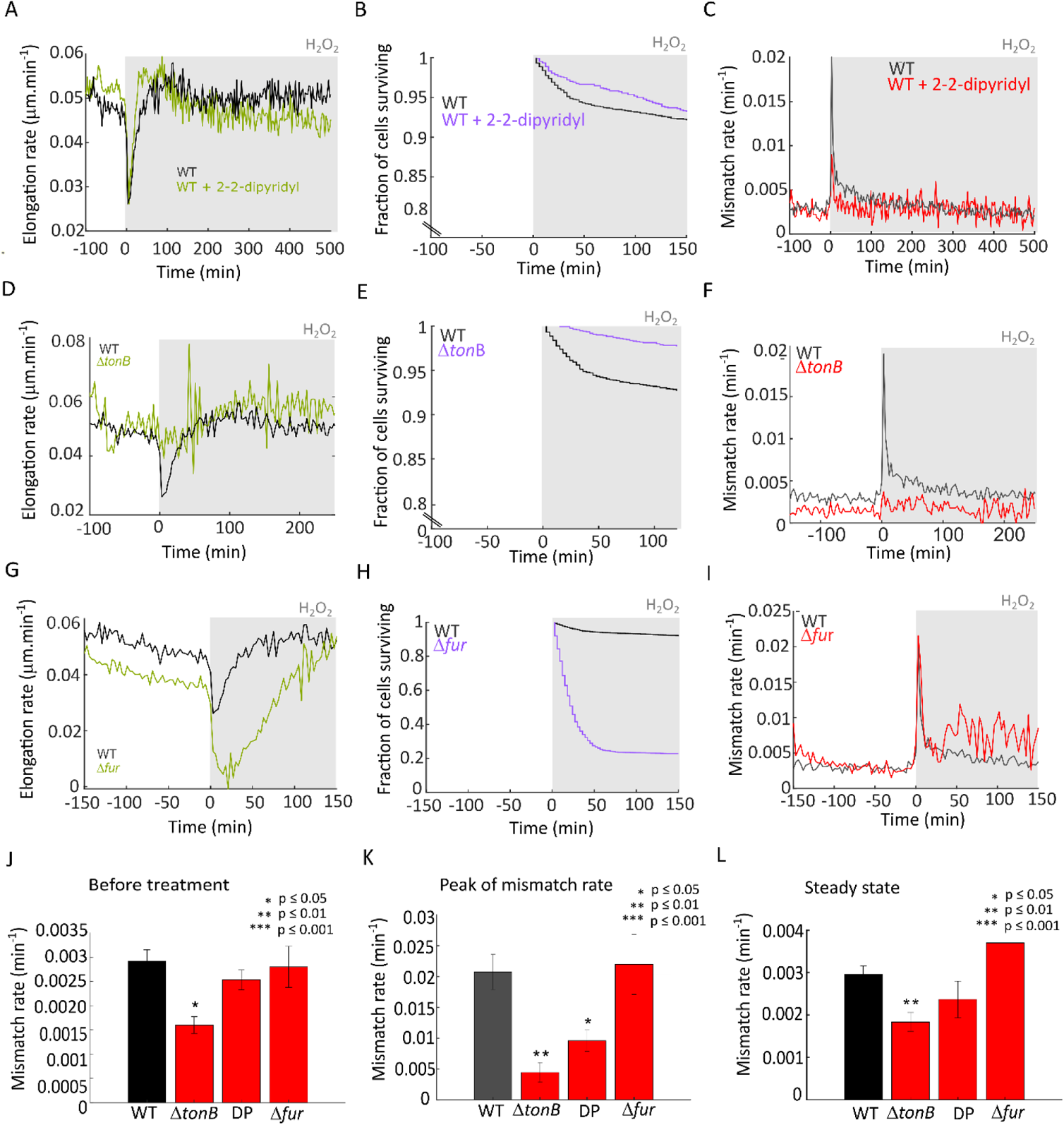
Perturbation of iron homeostasis impacts mutagenesis and adaptation to constant H_2_O_2_ treatment. (A-C) WT cells with 100 µM H_2_O_2_ treatment in medium supplemented with 50 µM 2-2-dipyridyl (DP, coloured lines) or without supplement (black). (A) Cell elongation rate (2081 cells, 3 experiments). (B) Fraction of cells surviving (1104 cells, 2 experiments). (C) Rate of DNA mismatches per cell per minute (2270 cells, 3 experiments). (D-F) Same as panels (A-C) for *tonB* deletion mutant (Δ*tonB*, coloured lines) and WT cells (black) with 100 µM H_2_O_2_ treatment. (D) Cell elongation rate (1523 cells, 3 experiments). (E) Fraction of cells surviving (1758 cells, 3 experiments). (F) Rate of DNA mismatches per cell per minute (1733 cells, 3 experiments). (G-I) Same as panels (A-C) for *fur* deletion mutant (Δ*fur*, coloured lines) and WT (black) treated with 100 µM H_2_O_2_. (G) Cell elongation rate (2593 cells, 4 experiments). (H) Fraction of cells surviving (3067 cells, 4 experiments). (I) Rate of DNA mismatches per cell per minute (2873 cells, 4 experiments). (J) Barplots showing the average mismatch rate per cell per minute for WT (black) and DP, Δ*tonB*, Δ*fur* cells (red) before treatment (mean ± SEM). Stars indicated the p-value of two-sample t-test. (K) Barplots showing the average mismatch rate per cell per minute for WT (black) and DP, Δ*tonB*, Δ*fur* cells (red), at the maximum of the mismatch rate peak with 100 μM H_2_O_2_ treatment (mean ± SEM). For Δ*tonB* the value was chosen as the average mismatch rate per cell per minute at the time corresponding to the average mismatch peak in WT cells. Stars indicated the p-value of two-sample t-test. (L) Barplots showing the average mismatch rate per cell per min for WT (black) and DP, Δ*tonB,* Δ*fur* cells (red) at the steady-state after the initial mutation burst during constant treatment with 100 μM H_2_O_2_ (mean ± SEM). Stars indicated the p-value of two-sample t-test.

## Discussion

ROS play a key role in mediating antibacterial functions of the immune system (Fang et al., 2016), antibiotic drugs (Giroux et al., 2017; Hong et al., 2019; Kohanski, Dwyer, et al., 2010), and during competition between bacterial species (Dong et al., 2015). The ability of bacteria to survive these stress conditions relies on oxidative stress responses, which induce conserved scavenging enzymes that provide effective resistance to ROS such as H_2_O_2_. Nevertheless, the sublethal concentrations of H_2_O_2_ that occur frequently in the environment (Imlay, 2019) are still highly mutagenic even in wild-type bacteria with functional oxidative stress responses (Bjelland & Seeberg, 2003; Rodríguez-Rojas et al., 2020), indicating an inherent vulnerability in the bacterial ROS defence mechanisms. Here, we set out to identify the nature of this weakness. Specifically, we investigated the chronology of H_2_O_2_-induced mutagenesis in relation to the timing of cell killing and the oxidative stress response in *E. coli*. We utilised microfluidic growth chambers to monitor intrinsic cellular dynamics while the environmental conditions and H_2_O_2_ concentrations were held constant.

We found that the OxyR response is very effective at preventing oxidative DNA damage and lethality, but the delay before the induction of H_2_O_2_ scavenging enzymes leaves cells vulnerable to the mutagenic effects of H_2_O_2_. Indeed, repeated H_2_O_2_ treatment leads to selection of mutants with a constitutively active OxyR response (Anand et al., 2020), which avoids the toxicity and mutagenesis caused by the adaptation delay (Fig. 4). In wild-type cells during the adaptation delay, DNA repair pathways are actively removing oxidative DNA damage, but a fraction of these lesions escapes repair and becomes fixed as mutations. Our findings suggest that such mutation dynamics are not unique to H_2_O_2_ stress. We previously showed that continuous alkylation stress also triggers a mutagenesis burst that is terminated when the expression of DNA repair enzymes is upregulated in cells (Uphoff, 2018). The mutagenesis bursts caused by oxidative stress and alkylation stress are fundamentally a consequence of adaptation delays, although the underlying molecular mechanisms are different. Whereas OxyR senses H_2_O_2_ within minutes and reliably activates stress response genes, the very low number of Ada proteins required for sensing alkylation damage delays the induction of DNA repair genes over multiple cell generations (Uphoff, 2018; Uphoff et al., 2013). Different from oxidative stress, cells adapted to alkylation stress still experience high levels of DNA damage, as shown by continuous elevated expression of the SOS response and sustained BER activity (Uphoff, 2018; Uphoff et al., 2013). However, once activated, these repair mechanisms are effective at removing mutagenic lesions such that the mutation burst is restricted to the period of the adaptation delay when the DNA repair capacity is low.

Our findings underline the notion that induction of reactive oxygen species as antibacterial agents is a double-edged sword. A brief period of accelerated mutation supply could speed up the evolution of antibiotic resistance and host adaptation. We speculate that a burst of mutations increases the chance of evolutionary rescue when an abrupt stress threatens a population to decline to extinction (Bell, 2017). However, it can take several generations before the expression of de novo mutations is sufficient to confer phenotypic stress resistance – a phenomenon known as phenotypic lag (Sun et al., 2018). In this case, stress responses can provide temporary tolerance for new mutants to survive the phenotypic lag. With live imaging, it is possible monitor mutation supply, stress responses, cell fitness, and population dynamics simultaneously. This approach complements genetic and population-scale studies to provide direct insights into evolutionary mechanisms.

## Materials and Methods

### Strains and plasmids construction

Unless otherwise indicated, strains were all derived from *Escherichia coli* AB1157 and constructed using classic molecular biology and genetics techniques. For microfluidic microscopy experiments, we used a previously constructed strain, carrying a Δ*flhD* gene deletion to inhibit cell motility, a constitutively-expressed fluorescent cell marker P_RNAI_-mKate2, and an endogenous MutL-mYPet fusion (Uphoff, 2018). It is known that *Escherichia coli* AB1157 carries an amber mutation in *rpoS* (Visick & Clarke, 1997), which may affect oxidative stress adaptation. We thus confirmed our central observations in separate experiments using *Escherichia coli* MG1655 as strain background (Fig. S17).

For single molecule tracking experiments, OxyR-Halo and Pol1-Halo fusions were constructed by lambda Red recombination (Datsenko & Wanner, 2000). We used a plasmid (pSU005, see plasmid list, (Banaz et al., 2019)) designed previously carrying an 11 amino acid linker (11 aa linker, SAGSAAGSGEF) followed by a HaloTag, and kanamycin resistance gene encompassed by two Flp recombinase recognition target (Frt) sites. To amplify these elements, we used forward primers containing 50 nt overhangs homologous to the C-terminal extremity of *oxyR*/*polA* gene and having the complementarity sequence to the 11aa linker. The reverse primers had 50 nt homology to the sequence immediately downstream of *oxyR*/*polA* gene (Table 3). We then inserted the PCR product on the chromosome using AB1157 cells containing the temperature-sensitive pkD46 plasmid which carries the lambda red recombination proteins controlled by an araBAD promoter. Cells grew in LB at 30°C to allow plasmid replication and 0.2 % arabinose was added to the medium to induce the expression of the recombination proteins. When cells reached 0.6 OD600, several cycles of centrifugation and resuspension of the pellet were performed using dH_2_O and 10 % glycerol. 14 µL of PCR product were then transformed by electroporation in 90 µL cells (2.5 kV electric pulse for 5 ms, Bio-Rad micropulser). Cells were recovered in SOC outgrowth medium (NEB B9020S) for 1 hour at 37°C before selection on kanamycin plates. pKD46 plasmid was cured by streaking single colonies twice at 37°C. Insertion was confirmed by colony PCR and the allele was moved to WT AB1157 by P1 phage transduction.

**Table 3:**
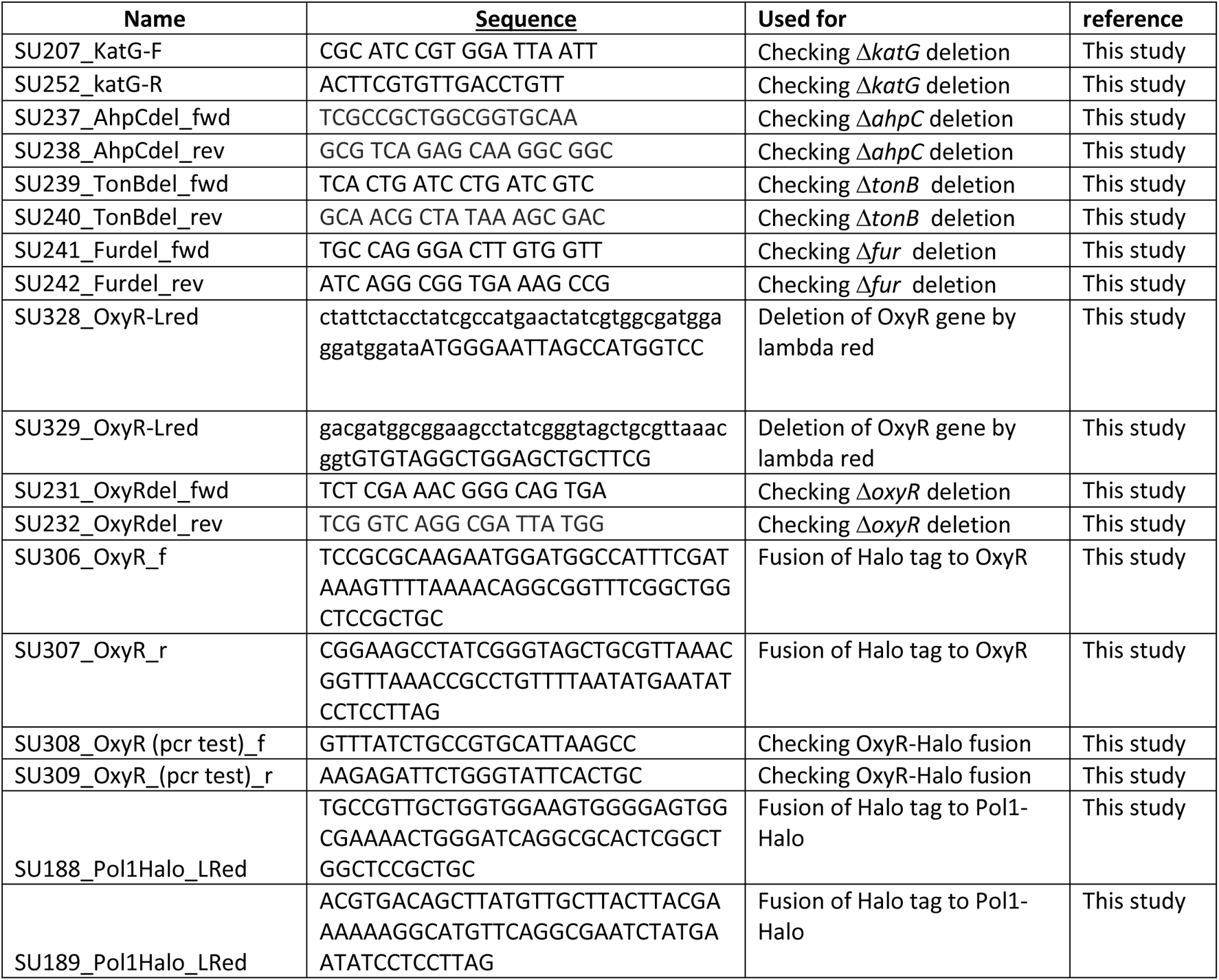
List of primers.

**Table 4.** List of reagents.

| Reagent | Number | company |
| --- | --- | --- |
| SOC medium | NEB B9020S | New England Biolabs |
| M9 minimal salts 5x | M9956 | Sigma |
| Solution 10x amino acids | 11130-036 | Gibco |
| L-Proline | A3453,0100 | Biochemica |
| Thiamine | A0955,0050 | Biochemica |
| Pluronic F-127 | P2443-250G | Sigma |
| Dow Corning Sylgard 184 kit | 1667370 | Farnell electronics |
| 1H,1H,2H,2H-perfluorooctyl silane | 448931 | Sigma |
| Coverslips No 1.5 24x50 mm | 631-0147 | VWR |
| Tygon Microbore silicon tubing | ND 100-80 | VWR |
| 30% W/W solution of H<sub>2</sub>O<sub>2</sub> | H1009-100mL | Sigma |
| 2,2- Dipyridyl | D216305 | Sigma |
| Catalase | C9322 | Sigma |
| TMR dye | G8251 | Promega |
| SDS-PAGE gels | XP00102 | Invitrogen |
| PageRuler Plus Prestained Protein Ladder 10 to 25 | 26619 | Thermo scientific |

All deletion mutants were moved to AB1157 by P1 transduction and antibiotic selection, using the corresponding Keio collection strains from the Coli Genetics Stock Center (CGSC) (Baba et al., 2006) Δ*katG*::kan (CGSC 10827), Δ*ahpC*::kan (CGSC 8713), Δ*tonB*::kan (CGSC 11229), Δ*fur::*kan (CGSC 8758), Δ*mutS*::kan (CGSC 10126). All gene deletions were checked by PCR using primers flanking the gene (Table 3). The antibiotic resistance flanked by frt sites was then removed using Flp recombinase expressed from pCP20 plasmid. We constructed the Δ*oxyR*∷kan mutant using lambda Red recombination and the plasmid pKD4 following the protocol from (Datsenko & Wanner, 2000) and as described above for the HaloTag fusions (Table 1).

**Table 1:** List of strains.

| Name | Genotype | Reference |
| --- | --- | --- |
| SU000 | AB1157 |  |
| SU471 | AB1157, PolA-11aa-Halo::kan | This study/(Banaz et al., 2019) |
| SU620 | AB1157, $\Delta flhD$, Tn7::mkate2, mutL-mYPet, carrying pUA066-PkatG-SCFP3A::kan (PSU033) | This study |
| SU621 | AB1157, $\Delta flhD$, Tn7::mKate2, mutL-mYPet, carrying pUA139 PrecA-SCFP3A::Kan (PSU032) | This study |
| SU699 | AB1157, OxyR-11aa-Halo::kan | This study |
| SU718 | AB1157, PolA-11aa-Halo, $\Delta fur$::kan | This study |
| SU777 | AB1157, $\Delta flhD$, Tn7::mKate2, mutL-mYPet, carrying pUA139 PgrxA-SCFP3A::Kan (PSU044) | This study |
| SU778 | AB1157, $\Delta flhD$, Tn7::mkate2, mutL-mYPet, $\Delta katG$ carrying pUA139-PgrxA-SCFP3A::kan (PSU044) | This study |
| SU779 | AB1157, $\Delta flhD$, Tn7::mKate2, mutL-mYPet, $\Delta ahpC$ carrying pUA139 PgrxA-SCFP3A::kan (PSU044) | This study |
| SU780 | AB1157, $\Delta flhD$, Tn7::mKate2, mutL-mYPet, $\Delta fur$ carrying pUA139 PgrxA-SCFP3A::kan (PSU044) | This study |
| SU802 | AB1157, $\Delta flhD$, Tn7::mkate2, mutL-mYPet, $\Delta oxyR$, carrying pUA139 PgrxA-SCFP3A::kan (PSU044) | This study |
| SU803 | AB1157, $\Delta flhD$, Tn7::mKate2, mutL-mYPet, $\Delta tonB$ carrying pUA139 PgrxA-SCFP3A::kan (PSU044) | This study |
| SU786 | AB1157, OxyR-Halo (SU701) + pUA139 PgrxA-SCFP3A kan (PSU044) | This study |
| SU887 | AB1157 pUA139-PgrxA-SCFP3A::kan (pSU044) | This study |
| SU888 | AB1157 $\Delta oxyR$- pUA139 PgrxA-SCFP3A::kan (pSU044) | This study |
| SU1072 | MG1655 $\Delta flhD$- Tn7::mkate2, mutL-mYPet + pUA139-PgrxA-SCFP3A Kan (PSU044) kan | This study |
| CM5255 | polA4113(ts) mutation | CGSC |

Reporter plasmids for the oxidative stress response were constructed based on pSC101 plasmids that expressed GFPmut2 under the control of PgrxA or PkatG (Zaslaver et al., 2006). The GFPmut2 was then replaced by the fast-maturing CFP variant SCFP3A (Balleza et al., 2018) using Gibson assembly. Similarly, we constructed a PrecA-SCFP3A reporter plasmid for the SOS response. All reporter plasmids were checked by sequencing and then inserted in different strains by heat-shock transformation (Table 2).

**Table 2:** List of Plasmids.

| Name | Characteristics |
| --- | --- |
| pSU044 | pUA139 PgrxA-SCFP3A kan |
| pSU033 | pUA066-PkatG-SCFP3A kan |
| pSU032 | pUA139-PrecA-SCFP3A kan |
| pSU005 | R6kgamma ori 11aa Halo frt kan frt |

### Microscopy

Before any microscopy experiment, strains were streaked from glycerol stocks stored at −80°C on LB plates with appropriate antibiotic and incubated overnight at 37°C.

#### Microfluidic microscopy

##### Cell culture before microscopy

For microfluidic microscopy, a single colony was grown in LB at 37°C with shaking for 6-8 hours, and the culture was diluted 1/50 overnight in 4 mL of M9 medium (M9 minimal salts 5x Sigma M9956) supplemented with 0.2% glucose, MEM amino acids (solution 10x Gibco 11130-036), 0.1 mg/mL L-proline (Biochemica A3453,0100), 0.5 μg/mL thiamine (Biochemica A0955,0050), 2mM MgSO_4,_ 0.1mM CaCl_2_ and the appropriate antibiotic. The next morning, overnight cultures were diluted 1/50 in 4 mL of the same M9 supplemented medium with antibiotic and incubated with shaking for 3 h at 37°C before loading in microfluidics devices. 0.85 mg/mL of surfactant pluronic F127 (Sigma, P2443-250G) was added to the morning culture to avoid cell aggregation in the device.

##### Preparation of the devices

Microfluidic experiments were performed with “mother machine” type of devices (P. Wang et al., 2010). Media is flowed in channels of sizes 100 µm x 25 µm (width x depth) and bacteria are in growth trenches of dimensions 1.2 µm x 1.2 µm x 20 µm (width x depth x length). These devices were prepared as described in (Moolman et al., 2013; Uphoff, 2018). Devices were made in Polydimethylsiloxane (PDMS) from a silicon wafer (produced by Kavli Nanolabs, Delft University) in two steps. An intermediate negative PDMS mold was made from the silicon wafer, by mixing monomer and curing agent 1:5 (Dow Corning Sylgard 184 kit (Farnell electronics, 1667370)). After removing bubbles in vacuum, the intermediate mold was cured at 65°C during 2 h, and subsequently treated overnight with Trichloro(1H,1H,2H,2H-perfluorooctyl)silane (Sigma 448931). The next day, after washing the intermediate mold with 100% ethanol, PDMS devices were made with 1:10 mix of monomer and curing agent. Bubbles were removed in vacuum and curing was done at 65°C for 2 h.

To make the preparation of the devices easier, we subsequently obtained silicon wafer from a different supplier (Conscience). In this case, the wafer had a negative relief, which means that PDMS devices could be made in one step (without an intermediate PDMS mold).

Before each experiment, one device was cut with a scalpel and holes for media inlet and outlet were inserted using a 0.75 mm biopsy punch. The device was then washed with 100% ethanol and dried with nitrogen gas 3 times. Microscope coverslips (No 1.5 24×50 mm, VWR 631-0147) were cleaned before bonding to the PDMS device. First coverslips were sonicated in acetone for 20 min, washed twice with dH_2_0, followed by 20 min of sonication in 100 % isopropanol and drying with nitrogen gas. Coverslips and devices were then bonded by treatment in air plasma (Plasma Etch PE-50) followed by 30 min in an oven at 95°C.

1 mL of the morning culture was concentrated by centrifugation and resuspended in 100 µL supernatant. Cells were then loaded in the PDMS device, by pipetting in the inlet/outlet holes. The whole device was then centrifuged for 10 min at 5000 rpm in a benchtop centrifuge with a custom holder to push the cells into the growth trenches. Supplemented M9 containing 0.85 mg/mL pluronic F127 without antibiotics was flowed via Tygon Microbore silicon tubing (VWR ND 100-80/0,508*1,524) that linked the device to a syringe pump (ALADDIN-220, World Precision Instruments) used with 30 mL syringes. Two syringes on two identical pumps were used for the experiment, both containing supplemented M9 and pluronic, and one of the two containing H_2_O_2_ (Sigma, 30% W/W solution H1009-100mL) at the concentrations stated in the text and figures. The tubing from the two syringes were linked to same device via a Y-junction with clamps to avoid backflow. During the first 10 min after loading, media was pumped at a rate of 2.5 mL/hr to flush out cells that were outside growth trenches and the flow rate was subsequently lowered to 0.5 mL/hr for data acquisition. Cells grew for 1 h in trenches before starting data acquisition to equilibrate to the environment. Data were acquired on average for 3 hours before switching to medium containing H_2_O_2_. Just after the start of the treatment the flow rate was increased for 10 min to 2.5 ml/hr to ensure rapid exchange of growth media and then lowered to 0.5 mL/hr for the remainder of the experiment. When indicated, 0,005 mg/mL 2,2’dipyridyl (DP, Sigma D216305, dissolved in DMSO) was added to supplemented M9 in both syringes (without and with H_2_O_2_). Catalase (Sigma C9322, 1 μg/mL) was added to the morning culture of the Δ*oxyR* mutant strain to help with growth and loading of the cells in the microfluidic chip. Catalase was only added in the culture until loading of the chip and not in the media used during data collection.

##### Imaging

Microfluidics experiments were performed on a Nikon Ti Eclipse inverted fluorescence microscope with a perfect focus system, oil immersion objective 100x NA1.4, motorized stage, sCMOS camera (Hamamatsu Flash 4), and LED excitation source (Lumencore Spectra X). The temperature chamber (Okolabs) allowed performing the experiments at 37°C. We recorded time-lapse movies with a frame rate of 1/3 min using NIS-Element software (Nikon) across three spectral channels (LED excitation wavelengths λ: 555 nm, 508 nm, 440 nm) to image mKate2, MutL-mYpet and CFP reporters respectively. One image is acquired from each channel consecutively using exposure times of 100 ms (λ = 555 nm), 300 ms (λ = 508 nm), and 75 ms (λ = 440 nm); LED intensity for all channels is set at 50% maximal output. One single triband dichroic and three separate emission filters were used to separate excitation and emission light.

Up to 48 fields of view can be imaged across the microfluidic device within the 3-minute time-lapse window. On average, 20 channels with bacteria were present per field of view such that about ~1000 mother cell traces were recorded per experiment.

##### Image Analysis

Time-lapse movies were analysed using custom scripts written in MATLAB (Mathworks). The shape of the mother cell at the closed end of each trench was automatically segmented based on the constitutive cytoplasmic mKate2 signal as described in (Norman et al., 2013). Cell length was calculated based on the segmentation mask and cell division events were identified as sharp drops in cell length. The generation time corresponds to the time interval between consecutive division events. The cell elongation rate was calculated from the difference in cell length per time interval. During H_2_O_2_ treatment, cell traces were truncated manually when cells ceased elongation and did not divide again until the end of the experiment. Filamentous cells escape from the growth trenches when their length reaches the open end. Cell traces of filamentous cells were manually truncated after their last division. For cells that lysed, traces were truncated automatically by the segmentation and lineage tracing algorithm. We quantified cell survival during H_2_O_2_ treatment as the fraction of cell traces before truncation relative to the number of cell traces at the start of the treatment. CFP reporter intensities were computed from the average CFP pixel intensities inside the segmentation mask area per cell, after subtracting the camera background signal outside of cells. To account for the fluorescence maturation time of SCFP3A at 37°C, we deconvoluted the observed intensity signal using an exponential function kernel with a decay constant of 6.4 min (Balleza et al., 2018). All intensity traces in the figures show the deconvoluted data. A spot-finding algorithm was used to detect MutL-mYPet foci, using a 3-pixel Gaussian band-pass filter and intensity thresholding. The cell-average mismatch rate per cell per minute corresponds to the total number of MutL-mYPet foci divided by the number of observed cells in each frame, and expressed in units of events/min based on the time interval of 3 min between frames. If a same mismatch event was detected in several consecutive frames, the event was counted only for the first frame. Plots of mismatch rate per cell per minute, CFP intensity, cell length, elongation rate, etc versus time show the average value at each time point across all cells, typically from 3 or more independent experiments performed on different days. Data from independent experiments were aligned temporally post-acquisition based on the time of H_2_O_2_ addition. Promoter activities of the gene expression reporters were computed from the difference in total CFP intensity per time interval, divided by the segmented cell area.

#### Functionality of HaloTag fusion proteins

The functionality of OxyR and Pol1 fusions to the HaloTag was tested by different assays detailed below.

##### H_2_O_2_ sensitivity assay

10-fold serial dilutions of the analysed strains were spotted on LB-agar plates containing no H_2_O_2_ or 100 μM H_2_O_2_. 2 sets of plates were made and incubated respectively at 30°C and 42°C as a control for the temperature-sensitive *polA*(ts) mutant strain (Fig. S11, Table 1).

##### Expression of OxyR-regulated fluorescent reporter

To assess the functionality of the OxyR-Halo fusion, we analysed the induction of the OxyR response using a plasmid carrying PgrxA-SCFP3A as a fluorescent transcriptional reporter. For consistency, we prepared cells from the WT AB1157 and OxyR-Halo fusion strains following the same procedure as for single-molecule tracking experiments (detailed below), but omitted the TMR dye to avoid cross-talk with the CFP reporter. Cells were treated by adding 100 μM H_2_O_2_ 40 min prior to data acquisition. Cells were immobilised on agarose gel pads (described below) and CFP intensities were measured on the Nikon TiE system (described above) using 50 % LED intensity at 100 ms exposure time. The CFP intensity per segmented cell area was obtained by segmentation of cell shapes from a phase contrast snapshot (described below) using MATLAB (Fig. S12).

##### Protein Gel

To confirm that HaloTag fusions are of the expected molecular weight and not degraded, we performed SDS PAGE in-gel fluorescence measurements. Strains were grown and labelled with TMR dye as for single-molecule tracking experiments, but with only one wash of medium to remove free dye (as opposed to 4 washes for single-molecule imaging). After labelling, the OD600 of the culture was adjusted to 0.05. 1 mL of culture was centrifuged, resuspended in 15 μL of 2X SDS loading buffer + 15 μL of dH_2_O, and incubated for 10 min at 95°C.

15 μL of sample was loaded on pre-casted SDS-PAGE gel (Invitrogen XP00102) with protein ladder (PageRuler Plus Prestained Protein Ladder 10 to 25, Life technologies 26619) and run for 1.5 h. The gel was imaged using 532 nm laser illumination (Fujifilm Typhoon FLA 7000) (Fig. S10).

#### Single molecule tracking

##### Cell cultures and labelling of the samples

For single-molecule imaging, a single colony was grown in LB with appropriate antibiotic for 6-8 h the day before the experiment. 8 µL of culture were diluted in 4 mL of supplemented M9 medium and incubated overnight with appropriate antibiotic. The next morning the culture was diluted 1/50 and incubated for 2 h with shaking at 37°C. 1 mL of culture was concentrated to 100 μL before labelling the Halo-tag as described in (Banaz et al., 2019). Briefly, 5 μL of 50 μM stock solution of TMR dye (Promega G8251) was added to the concentrated culture, mixed vigorously, and incubated for 30 min at 25°C. The labelled culture was centrifuged and washed 4 times with 1 mL supplemented M9 medium to remove free dye. Cells were then recovered at 37°C with shaking for 30 min, washed again, and 1 μL of concentrated cell suspension was pipetted on an agarose pad (1% agarose in supplemented M9 medium, with or without 100 µM H_2_O_2_) and sandwiched between two coverslips. Imaging was started as soon as possible after cells were placed on the pads (within ~5 minutes).

##### Single-molecule tracking

Single-molecule tracking was performed using a custom-build Total Internal Reflection Fluorescence Microscope (TIRF) under oblique illumination (Banaz et al., 2019; Wegel et al., 2016) and controlled via Andor Solis software. Movies of 10.000 frames with 15.48 ms time interval between frames were acquired under constant 561 nm excitation laser illumination at 0.5 kW/cm^2^. Each field of view was exposed to 561 nm constant laser illumination at 0.5 kW/cm^2^ for about 30 seconds before acquisition to deactivate the majority of TMR fluorophores for single-molecule detection and tracking. The acquisition time of each movie was noted relative to the time since the cells had been placed on the agarose pad with or without H_2_O_2_ treatment. An image of each field of view was also acquired with transmitted light from LED illumination for cell segmentation. All single-molecule tracking experiments were performed at room temperature and were repeated at least three times on different days for each condition.

Single-molecule tracking data were analysed in MATLAB as previously described (Stracy et al., 2016). The mean diffusion coefficient (D) for each track was calculated from the Mean Squared Displacement over 5 frames (4 steps). The distribution of diffusion coefficients was fitted with an analytical model of two species of molecules with distinct diffusion coefficients. Free fit parameters are the mean diffusion coefficient of each species and their relative abundance. Fitting the combined data from all measurement experiments and time points yielded a mean diffusion coefficient of the immobile population of molecules of 0.17 µm^2^/s. This value was subsequently constrained for fitting the distributions of each time point separately. The diffusion coefficient of the second population of mobile molecules was left unconstrained.

### Rifampicin assay

Mutation frequency was measured using rifampicin resistance assays following the protocol from (Rodríguez-Rojas et al., 2020) and adapting it to the conditions of this study (Fig. S7). Briefly, 3 cultures of WT cells were grown until OD ~0.2 in LB medium. At this time, the volume of each culture was increased to 50 mL and then divided in 2 tubes of 25 mL. The former was kept untreated and the latter was treated with 1 mM H_2_O_2_ and 5 mL of culture were collected at different treatment times. Treatment was stopped at 5, 12, 20, 27 and 35 minutes by adding 10 μg/mL of catalase (Sigma C9322) to the 5 mL of culture and by then centrifuging for 6 min at 4000 rpm. Treated and untreated cultures were then concentrated in 1 mL LB and grown at 37°C overnight. The next morning, 100 µL of culture was plated on LB plates containing 100 μg/mL of rifampicin. Serial dilutions were performed and 100 µL of a 10^−7^ dilution of each culture were plated on LB plates without rifampicin. We calculated mutation frequencies by dividing the number of colonies counted on LB plates containing rifampicin by the number of colonies on LB plates without rifampicin for each culture. For each condition, the experiment was performed on three separate days with three independent cultures per day. Mean mutant frequency and standard error of the mean for the 9 total cultures were calculated and two-sample t-test performed.

## Acknowledgements

We thank all members of the Uphoff lab and former members of David Sherratt’s lab for discussions. Research in the Uphoff lab is funded by a Wellcome Trust & Royal Society Sir Henry Dale Fellowship (206159/Z/17/Z), a WellcomeBeit Prize (206159/Z/17/B) and a Research Prize Fellowship of the Lister Institute of Preventative Medicine.

## Author contributions

### Valentine Lagage

Department of Biochemistry, University of Oxford, Oxford, United Kingdom

Contribution : conceptualization – methodology – software-validation - formal Analysis – investigation – data Curation - Writing – Original Draft Preparation - Writing – Review & Editing – Visualization – Supervision - Project Administration

Conflict of interest – No conflict of interest declared

### Victor Chen

Department of Biochemistry, University of Oxford, Oxford, United Kingdom

Contribution: methodology - software-validation - formal Analysis – investigation - data Curation Conflict of interest – No conflict of interest declared

### Stephan Uphoff

Department of Biochemistry, University of Oxford, Oxford, United Kingdom

Contribution: conceptualization – methodology – software-validation - formal Analysis – resources - data Curation - Writing – Original Draft Preparation - Writing – Review & Editing – Visualization – Supervision-Project Administration - Funding Acquisition

Conflict of interest – No conflict of interest declared

## Supplementary figures

**Fig. S1:**
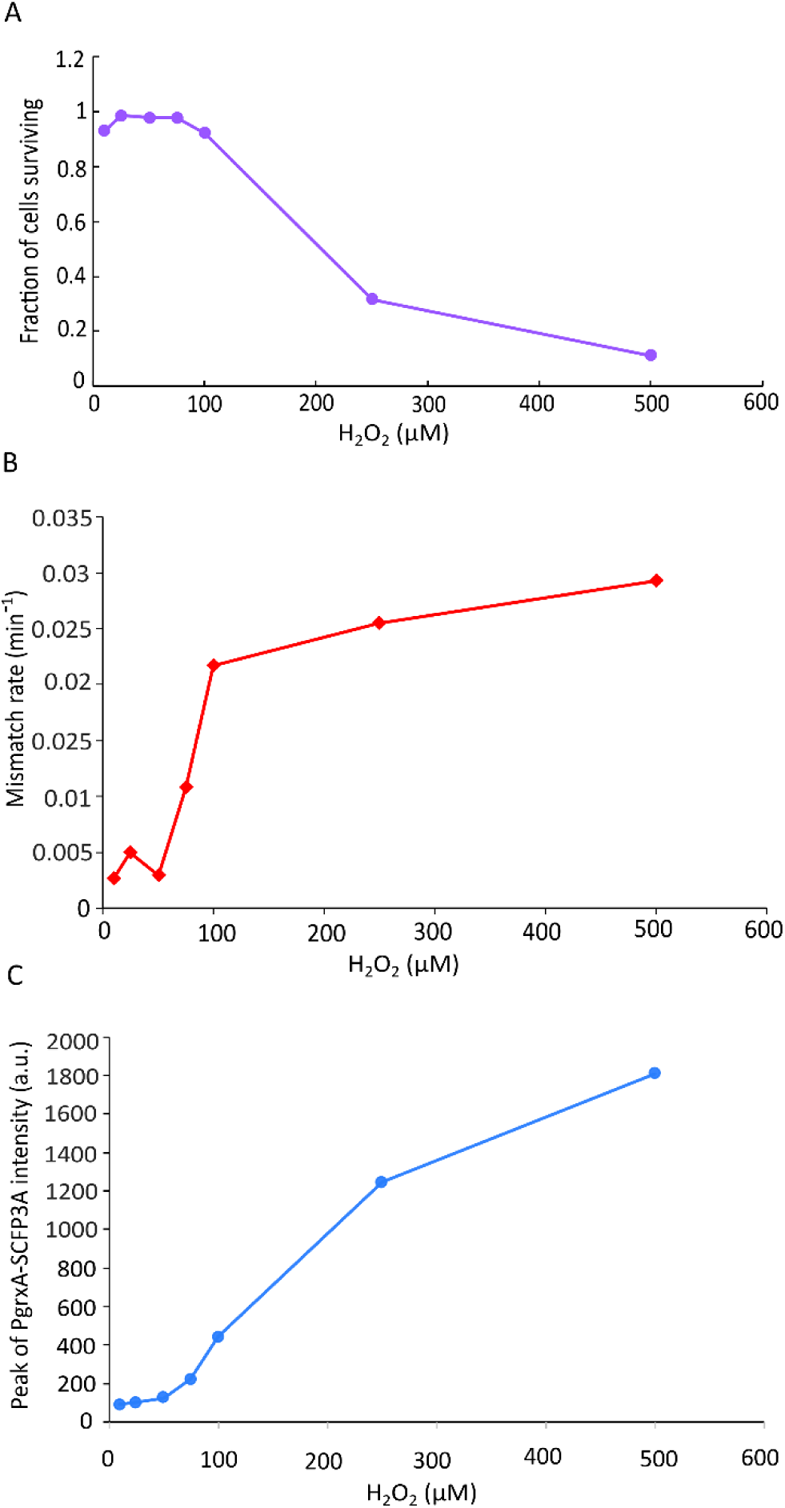
Survival after 100 min, mismatch rate peak, and peak expression of the oxidative stress response in cells treated with different doses of H_2_O_2_. (A) Fraction of cells surviving after 100 minutes of treatment with different doses of H_2_O_2_ (10 μM (1 experiment 656 cells), 25 μM (2 experiments, 1314 cells), 50 μM (476 cells), 75 μM (1 experiment, 471 cells), 100 μM (8 experiments, 6657 cells), 250 μM (1 experiment, 335 cells), 500 μM (1 experiment, 772 cells)). (B) Rate of DNA mismatches per cell per minute at the peak of the mutagenesis burst for different doses of H_2_O_2_ (10 μM (1 experiment, 651 cells), 25 μM (1 experiment, 605 cells), 50 μM (1 experiment, 465 cells), 75 μM (1 experiment, 466 cells), 100 μM (8 experiments, 6655 cells), 250 μM (1 experiment, 331 cells), 500 μM (1 experiment, 764 cells)). (C) Peak of PgrxA-SCFP3A reporter intensity in response to different doses of H_2_O_2_ (10 μM (1 experiment, 651 cells), 25 μM (2 experiments, 1281 cells), 50 μM (1 experiment, 465 cells), 75 μM (1 experiments, 466 cells), 100 μM (2 experiments, 1848 cells), 250 μM (1 experiment, 331 cells), 500 μM (1 experiment, 764 cells)).

**Fig. S2:**
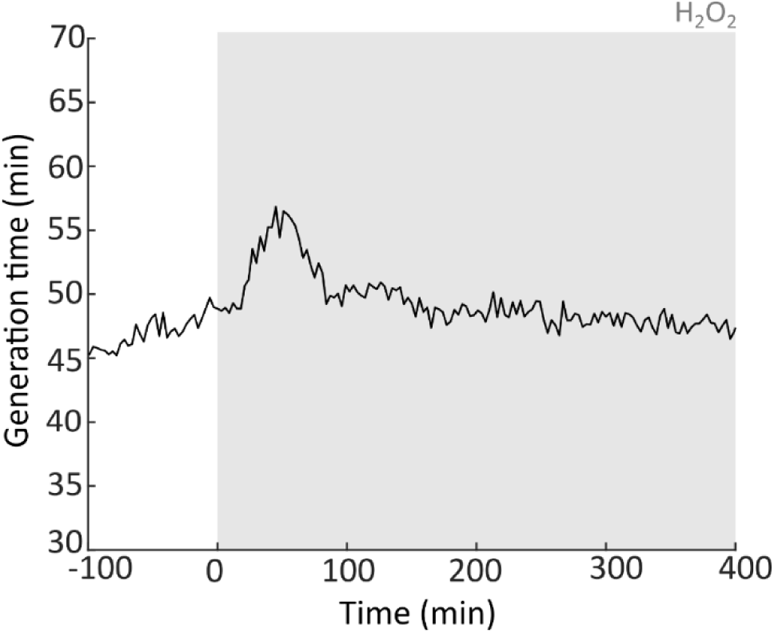
Transient increase of the generation time in WT cells treated with H_2_O_2_. Generation time of WT cells treated with 100 μM of H_2_O_2_. (8 experiments, 6562 cells).

**Fig. S3:**
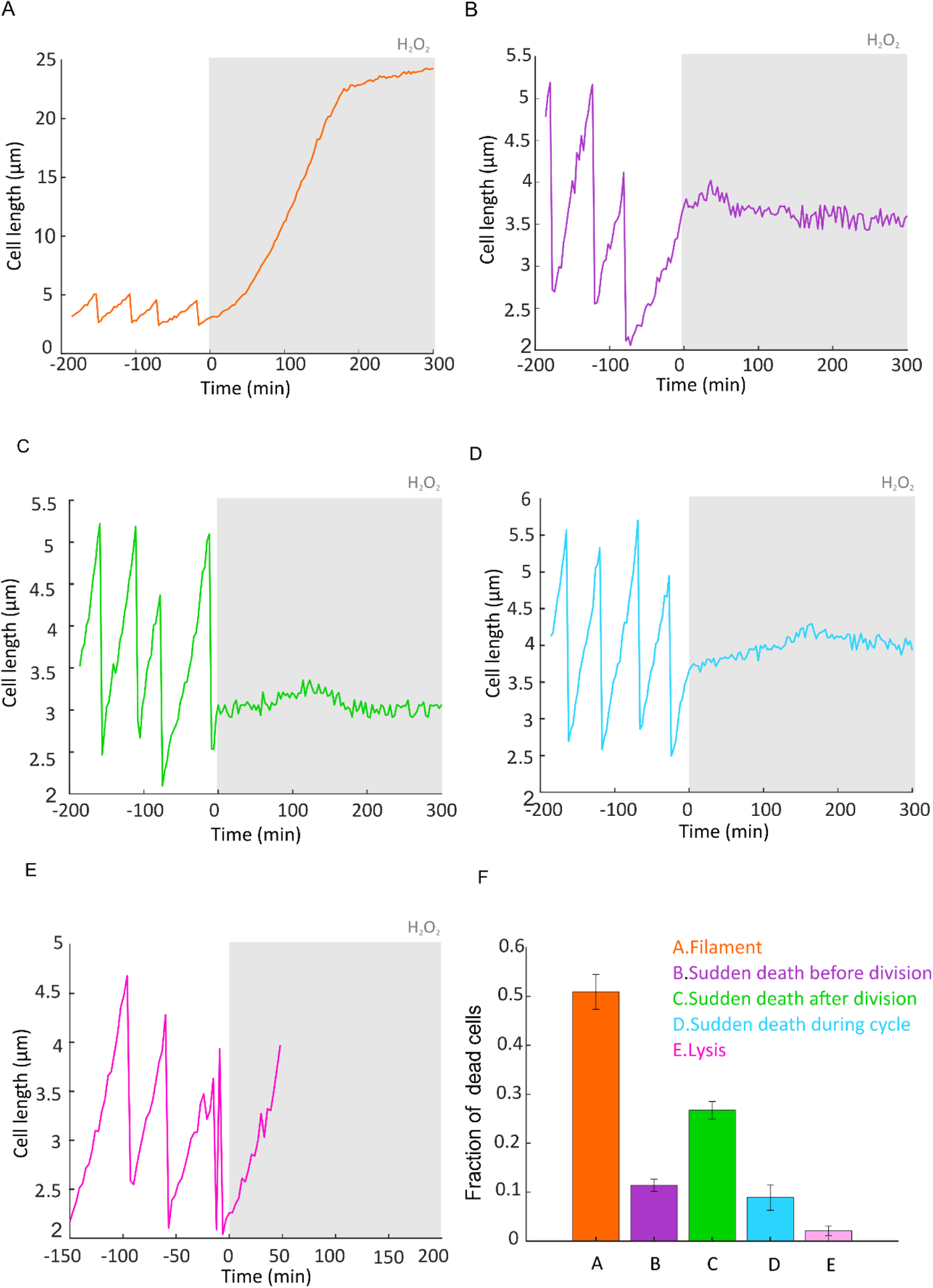
Cell length traces show different types of cell death with 100 µM H_2_O_2_ treatment. (A) Example cell length trace showing filamentation. (B) Example cell length trace showing growth arrest just before cell division. (C) Example cell length trace showing growth arrest just after cell division. (D) Example cell length trace showing growth arrest during the cell cycle. (E) Example cell length trace showing lysis. (F) Percentage of cells for each type of cell death (6562 cells, 8 experiments, mean ± SEM).

**Fig. S4:**
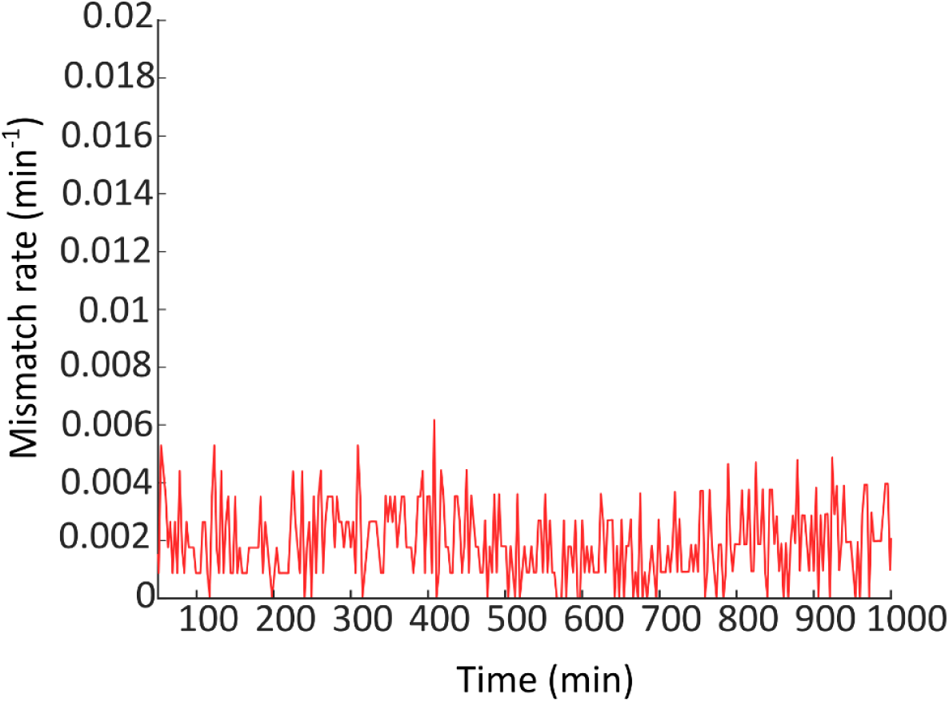
Mismatch rate is stable over time in untreated cells. Rate of DNA mismatches per cell per minute in untreated cells (651 cells, 1 experiment).

**Fig. S5:**
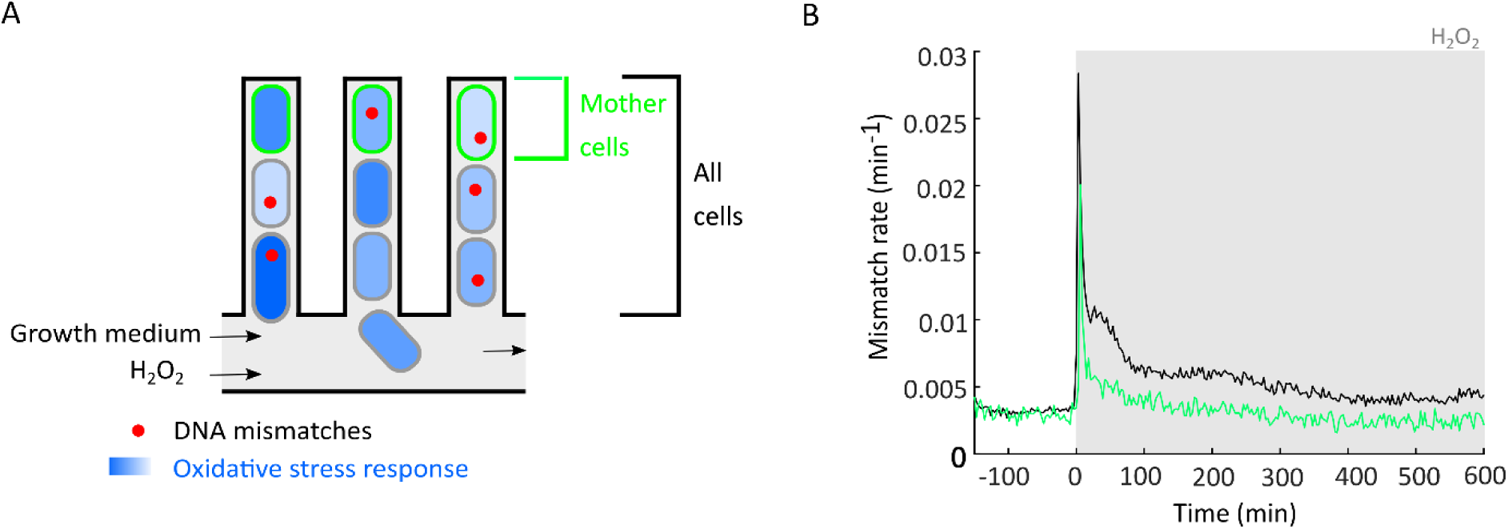
All cells in the microfluidic growth trenches experience a burst of mutagenesis. (A) Throughout the study, we focused our analysis on the mother cells that can be continuously observed at the closed end of each growth trench over multiple generations. (B) To confirm that the burst mutagenesis is not limited to these cells, we quantified the rate of DNA mismatches per cell per minute for all cells in the trenches with 100 µM H_2_O_2_ treatment (35082 cells, 8 experiments).

**Fig. S6:**
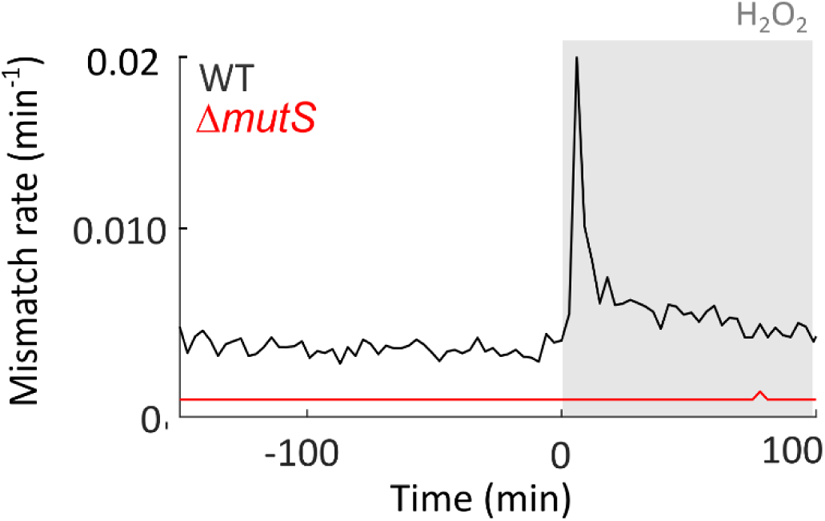
No MutL-mYPet foci are detected in a Δ*mutS* deletion strain. (A) Rate of DNA mismatches per cell per minute of a Δ*mutS* mutant strain with 100 µM H_2_O_2_ treatment (551 cells, 1 experiment).

**Fig. S7:**
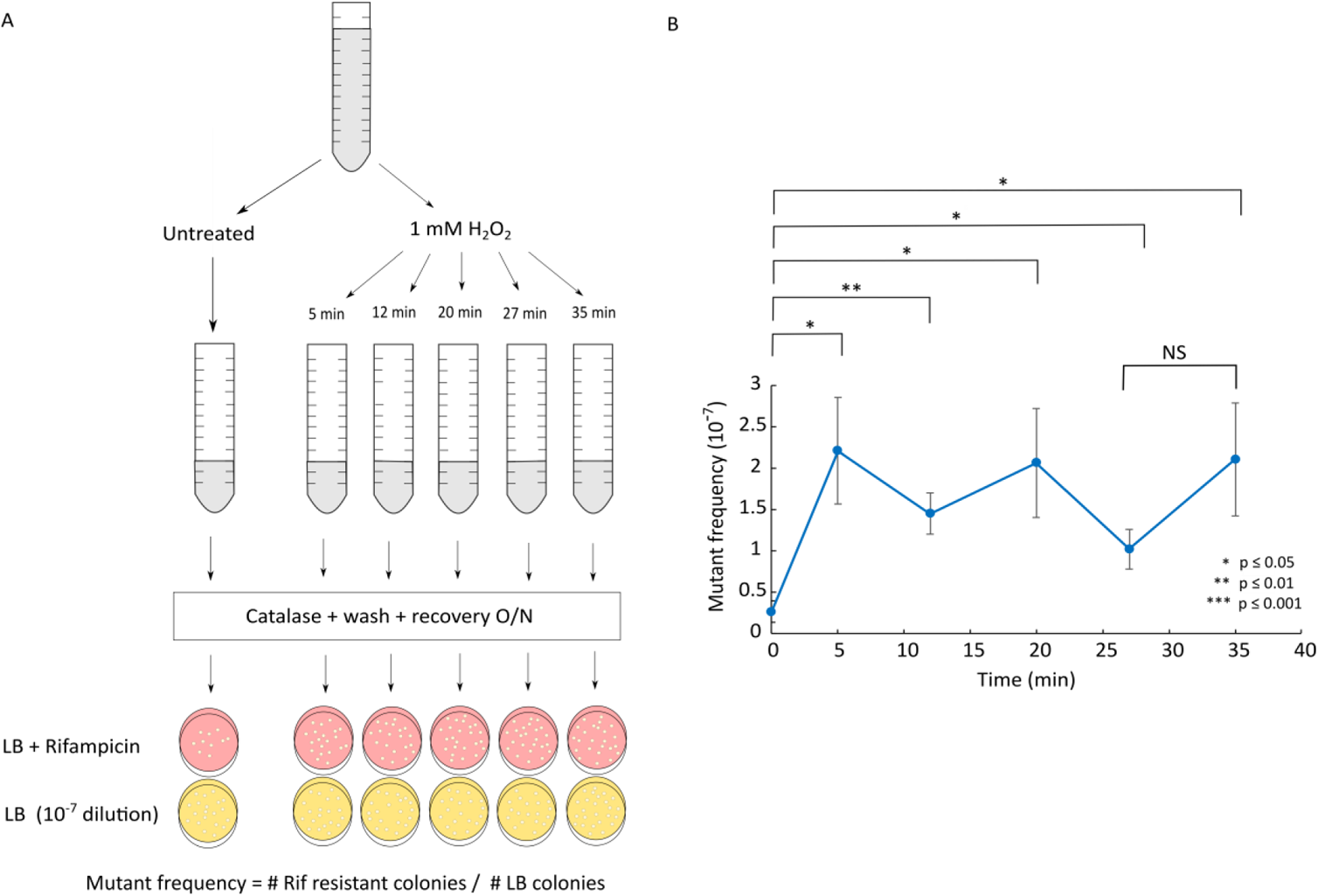
Measurement of the genomic mutation frequency confirms that H_2_O_2_ causes a mutagenesis burst at the start of treatment. **(A)** Cultures were treated with 1 mM H_2_O_2_ for 5, 12, 20, 27 and 32 min by adding catalase, then washed with M9 without H_2_O_2_ and recovered overnight before plating on LB + Rifampicin and a 10^−7^ dilution of each culture was plated on LB. Mutant frequency was quantified from the ratio of Rifampicin-resistant colonies divided by the colony count on LB plates. **(B)** The frequency of rifampicin-resistant colonies (a reporter for mutation frequency, mean ± SEM, 6-9 experiments) increases after 5 minutes of 1 mM H_2_O_2_ treatment but does not increase further during prolonged treatment. Stars or NS (not significant) show the p-value of two-sample t-tests.

**Fig. S8:**
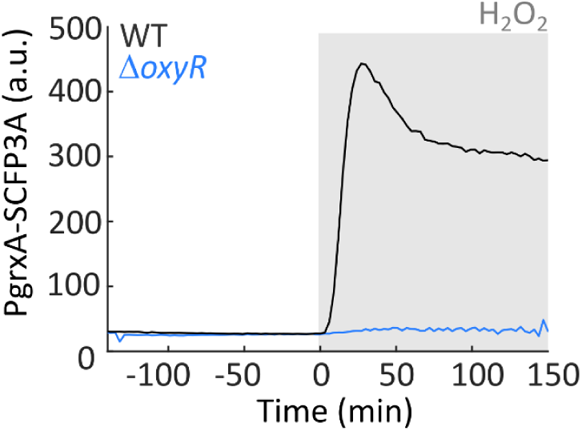
PGrxA expression is not induced in Δ*oxyR* mutant cells. PgrxA-SCFP3A expression stays at a basal level in Δ*oxyR* cells treated with 100 μM H_2_O_2_ (728 cells, 3 experiments), compared to WT (black).

**Fig. S9:**
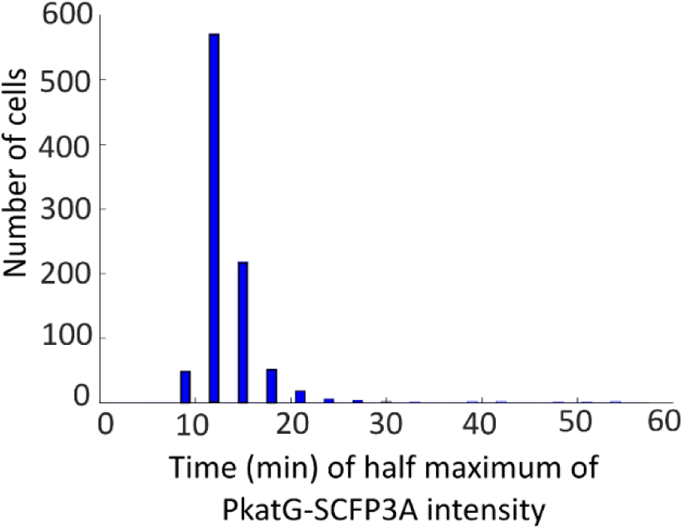
KatG expression is induced shortly after the start of the treatment. Histogram of the lag time distribution for PkatG-SCFP3A to reach half-maximal intensity after start of 100 µM H_2_O_2_ treatment (988 cells, 1 experiment).

**Fig. S10:**
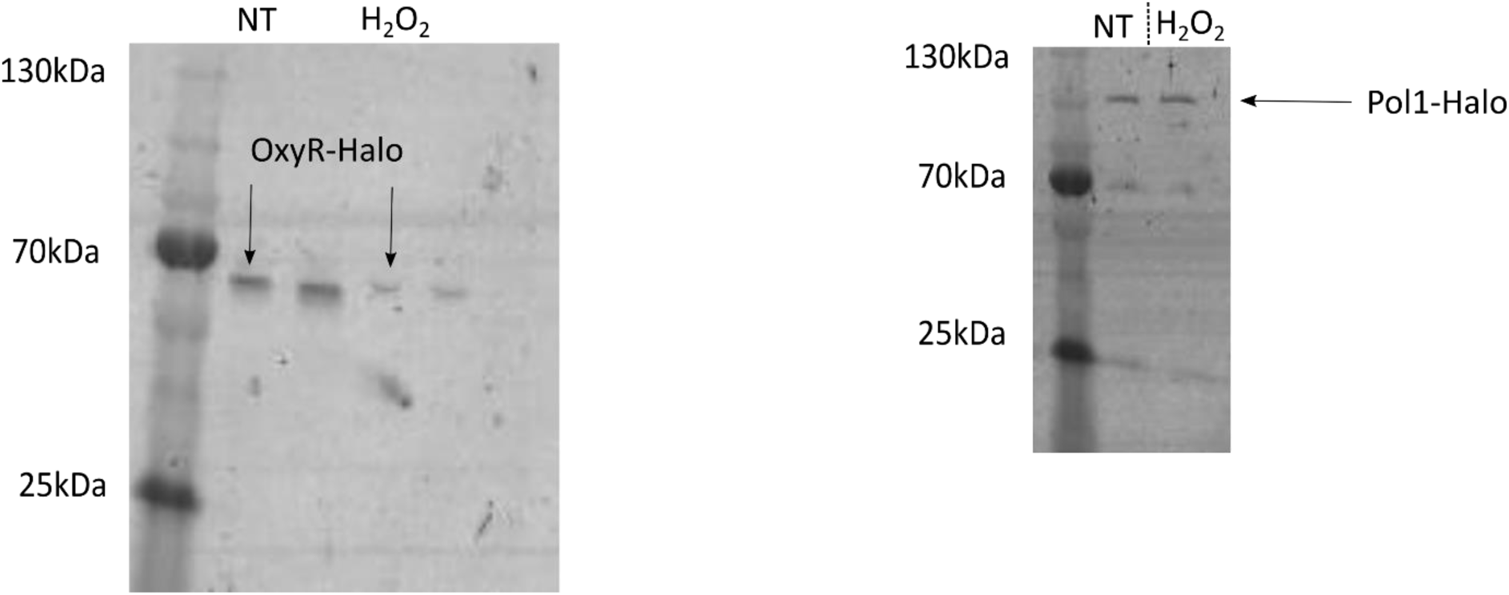
In-gel fluorescence confirms stability of OxyR-Halo and Pol1-Halo fusions. Strains expressing OxyR-Halo and Pol1-Halo were labelled with TMR dye as for microscopy experiments. Cells were lysed and analysed via SDS-PAGE without treatment (NT) and 30 min after treatment with 100 μM H_2_O_2_. For OxyR-Halo fusion, one band is detected on the gel corresponding to the expected size of OxyR (34.4 kDa) plus the HaloTag (33 kDa). For Pol1-Halo fusion, one band is detected on the gel corresponding to the expected size of Pol1 (94.08 kDa) plus the HaloTag (33 kDa). The 2 unlabelled lanes in the OxyR-Halo gel are not relevant to this work.

**Fig. S11:**
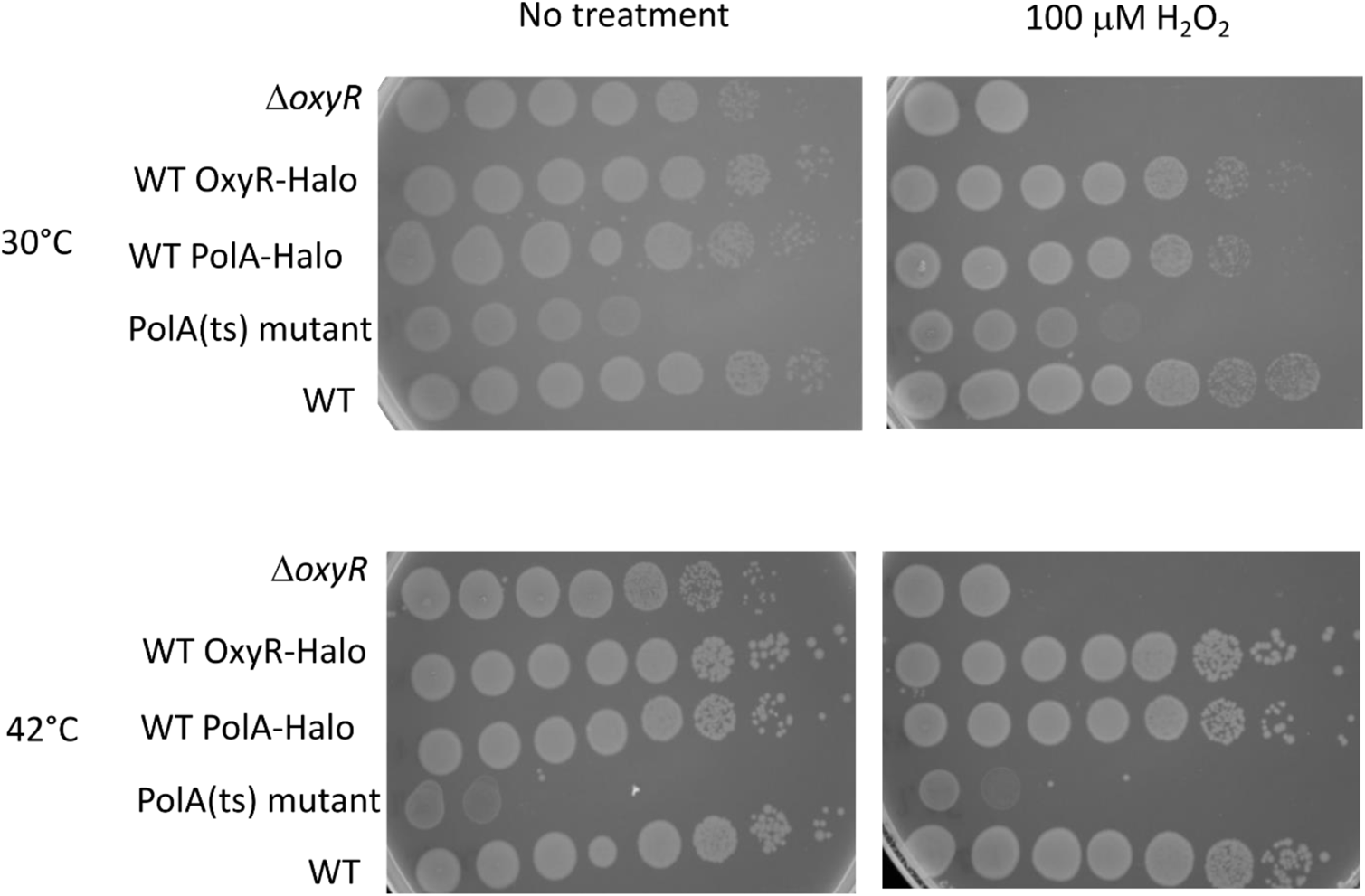
Survival assay to assess the functionality of OxyR-Halo and Pol1-Halo fusions. Survival assay without H_2_O_2_ treatment and with 100 µM H_2_O_2_ treatment performed at 30°C and at 42°C as PolA(ts) mutant is thermosensitive.

**Fig. S12:**
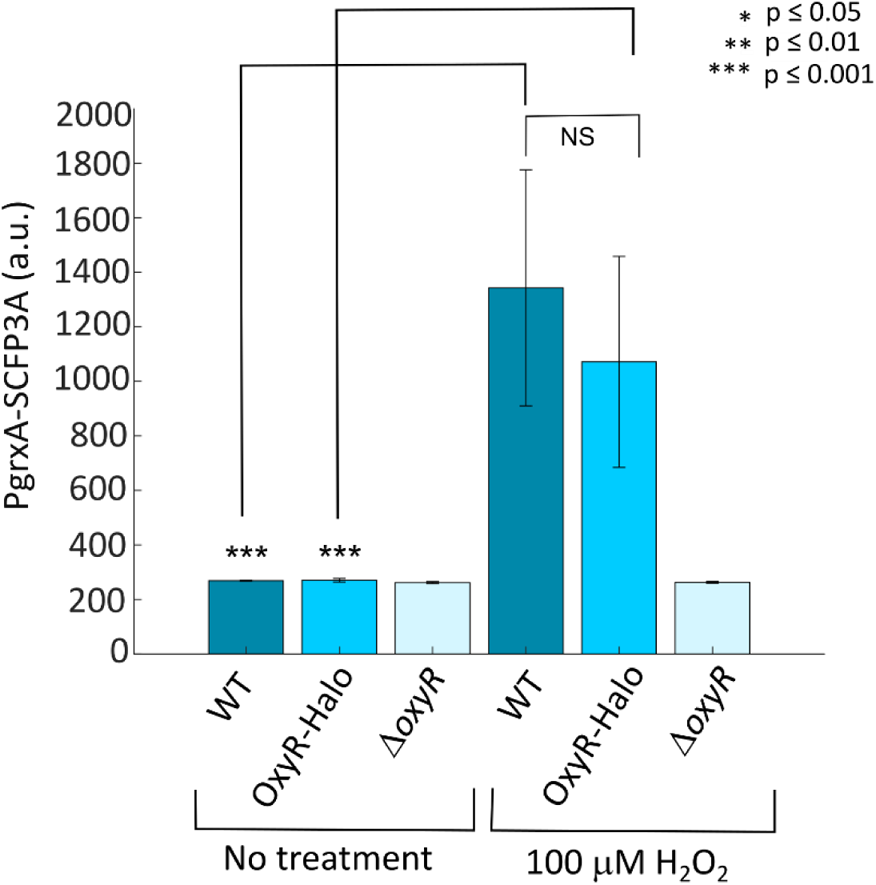
Induction of OxyR-dependent gene expression confirms functionality of the OxyR-Halo fusion. Barplots of the average PgrxA-SCFP3A intensity in WT, OxyR-Halo and Δ*oxyR* mutant cells without treatment (mean ± SEM, 4 experiments, 3748 cells for WT, 3682 cells for WT OxyR-Halo, 2266 cells for Δ*oxyR*), and with 100 µM H_2_O_2_ treatment (mean ± SEM, 4 experiments, 3106 cells for WT, 4336 cells for WT OxyR-Halo, 2816 cells for Δ*oxyR*. P-values of two-sample t-tests are indicated or NS (not significant)

**Fig. S13:**
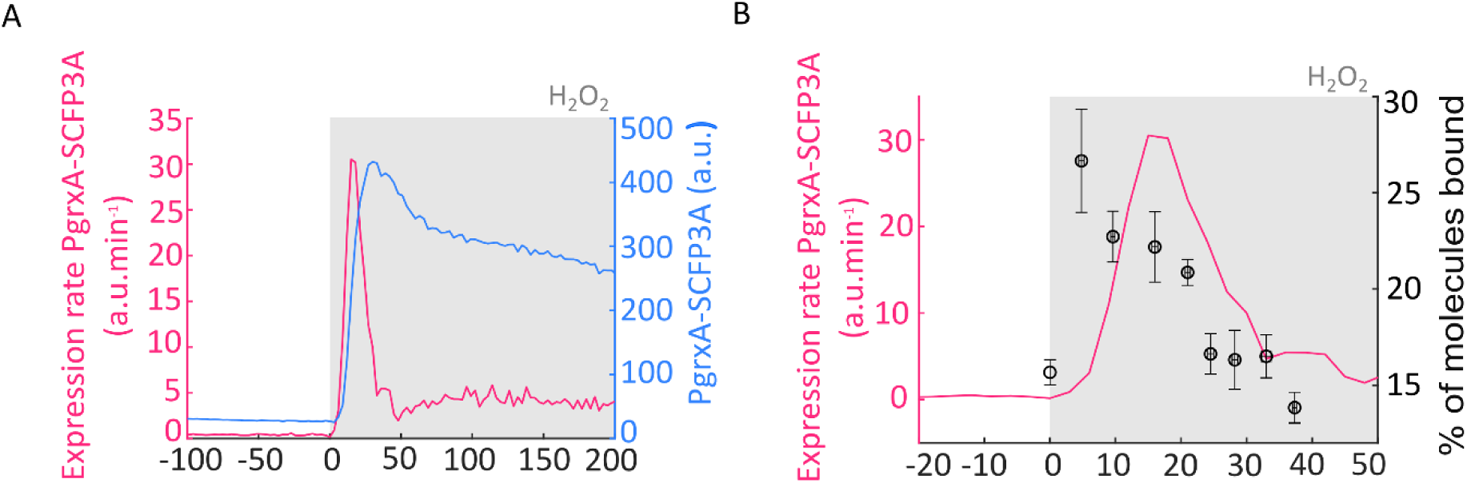
Expression rate and intensity of PgrxA-mSCFP3 reporter. (A) Expression rate of PgrxA-SCFP3A (pink, 878 cells, 1 experiment) and PgrxA-SCFP3A intensity (blue, 878 cells, 1 experiment) with 100 µM H_2_O_2_ treatment. (B) Expression rate of PgrxA-SCFP3A (pink, 878 cells, 1 experiment) and percentage of bound OxyR-Halo molecules (black, from Fig. 3D).

**Fig. S14:**
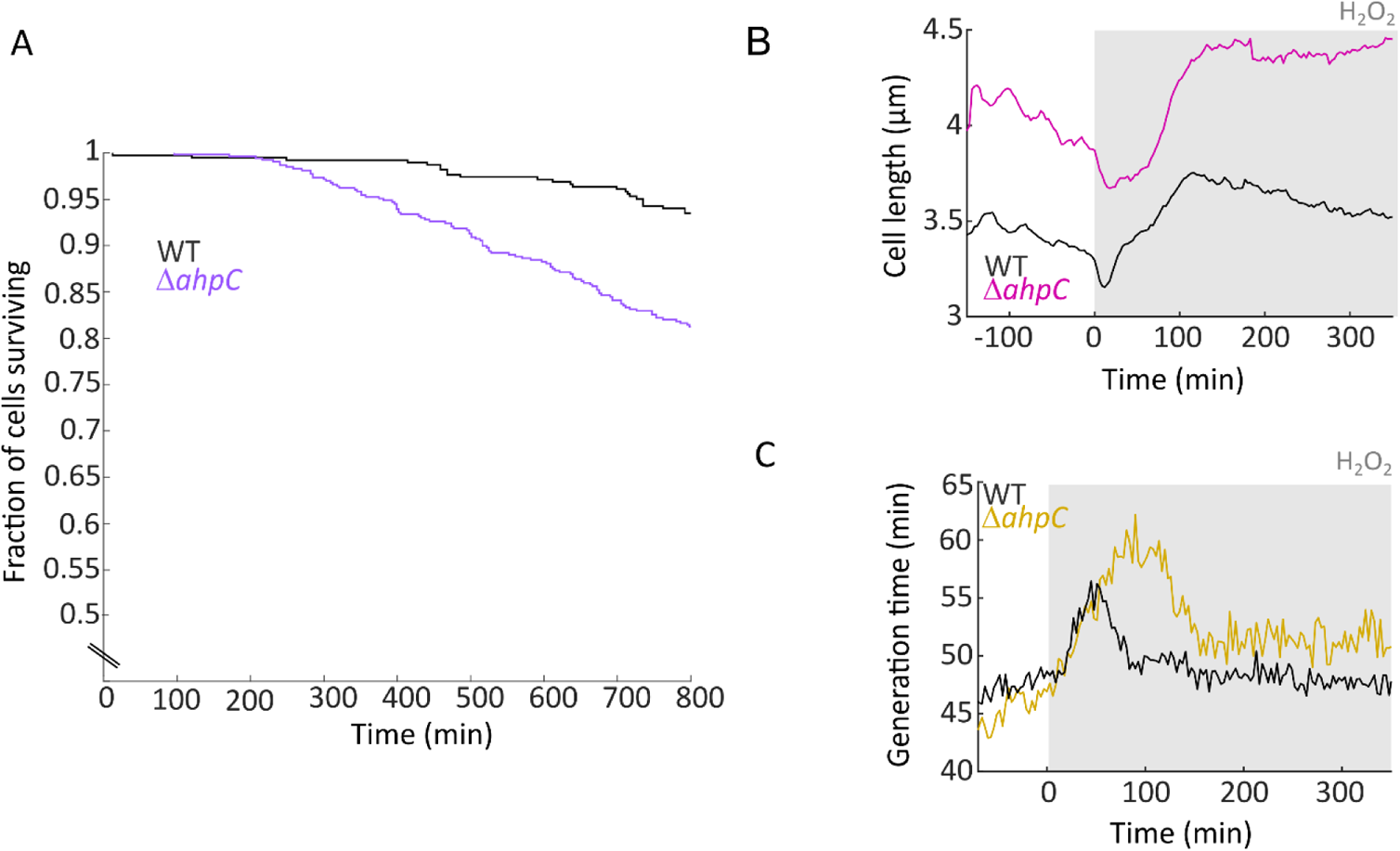
Growth characteristics of Δ*ahpC* mutant cells compared to WT cells. Fraction of cells surviving without treatment for WT cells (black) and Δ*ahpC* mutant cells (purple, 528 cells). The elongation rate of the Δ*ahpC* mutant is higher than for WT cells before and during 100 µM H_2_O_2_ treatment (Fig. 4D). This can be explained by its higher cell length (B, pink, 3360 cells, 5 experiments) compared to WT (black) while the generation time (A, purple, 3376 cells, 5 experiments) is unchanged before treatment compared to WT, but increased during 100 µM H_2_O_2_ treatment (black).

**Fig. S15:**
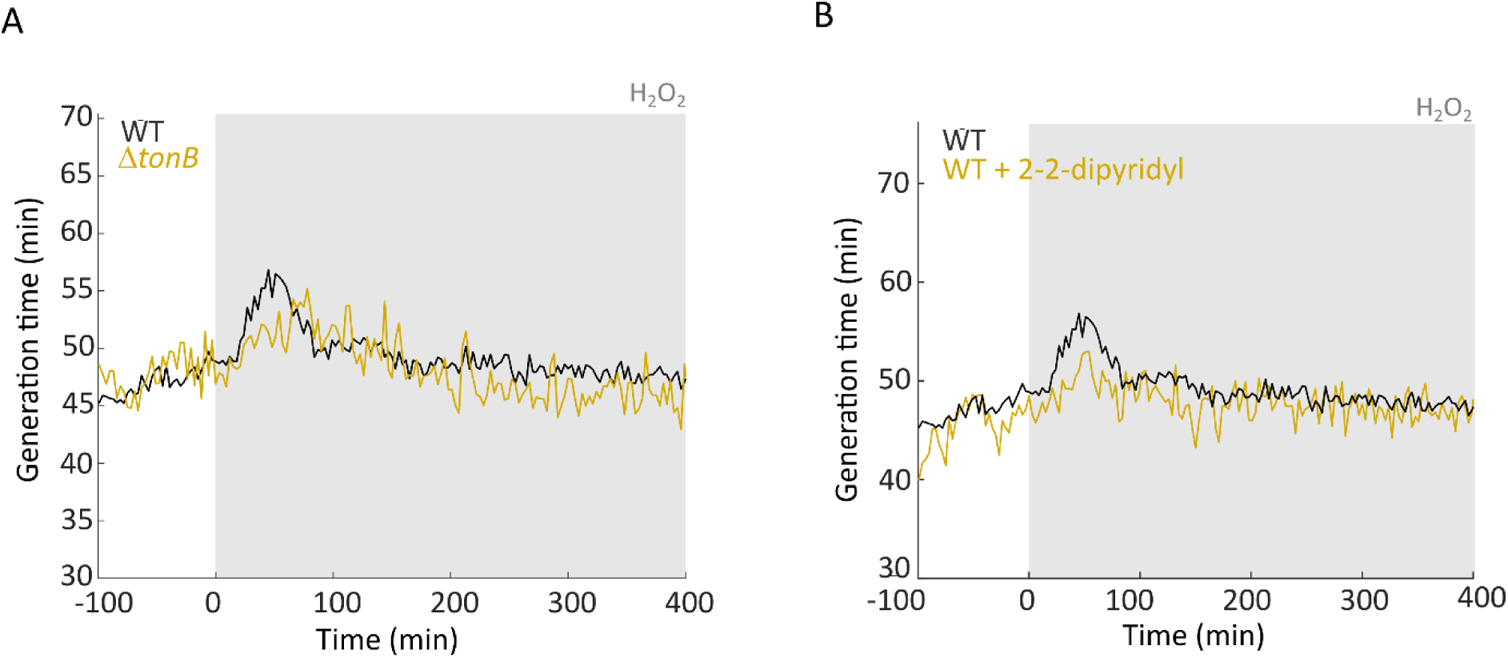
Effect of iron limitation on cell cycle duration with H_2_O_2_ treatment. (A) Generation time of WT (black) and Δ*tonB* (yellow, 1753 cells, 3 experiments) mutant cells with 100 µM H_2_O_2_ treatment. (B) Generation time of WT cells with 100 µM H_2_O_2_ treatment in medium supplemented with 50 µM 2-2-dipyridyl (yellow, 2010 cells, 3 experiments) or without supplement (black).

**Fig. S16:**
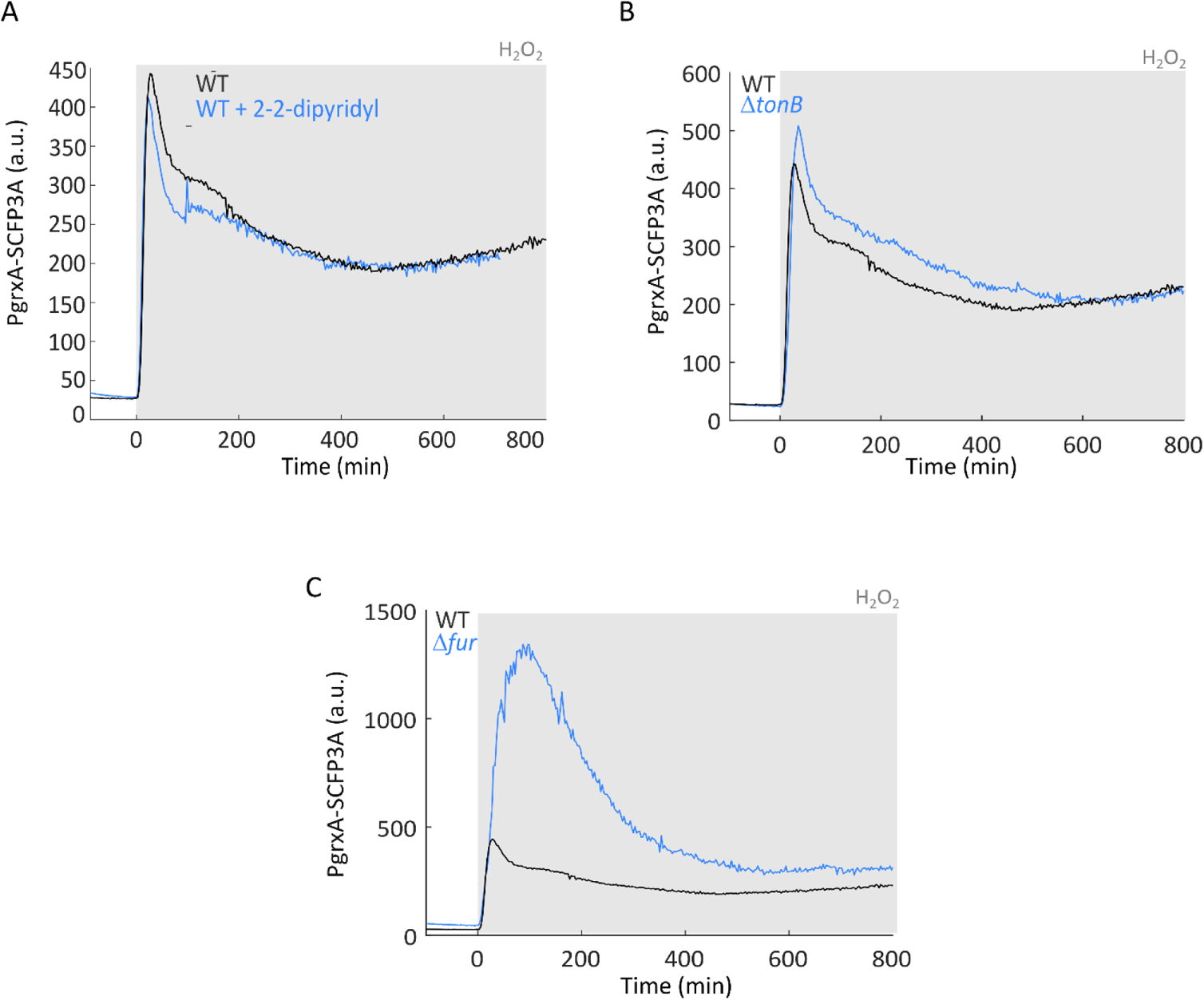
Effects of perturbed iron homeostasis on OxyR response expression with 100 µM H_2_O_2_ treatment. (A) PgrxA-SCFP3A intensity of cells in presence of DP (2270, 3 experiments cells). (B) PgrxA-SCFP3A intensity of Δ*tonB* mutant (573 cells, 2 experiments). (C) PgrxA-SCFP3A intensity of Δ*fur* mutant (520 cells, 1 experiment). WT is shown in black for comparison (2355 cells, 2 experiments).

**Fig. S17:**
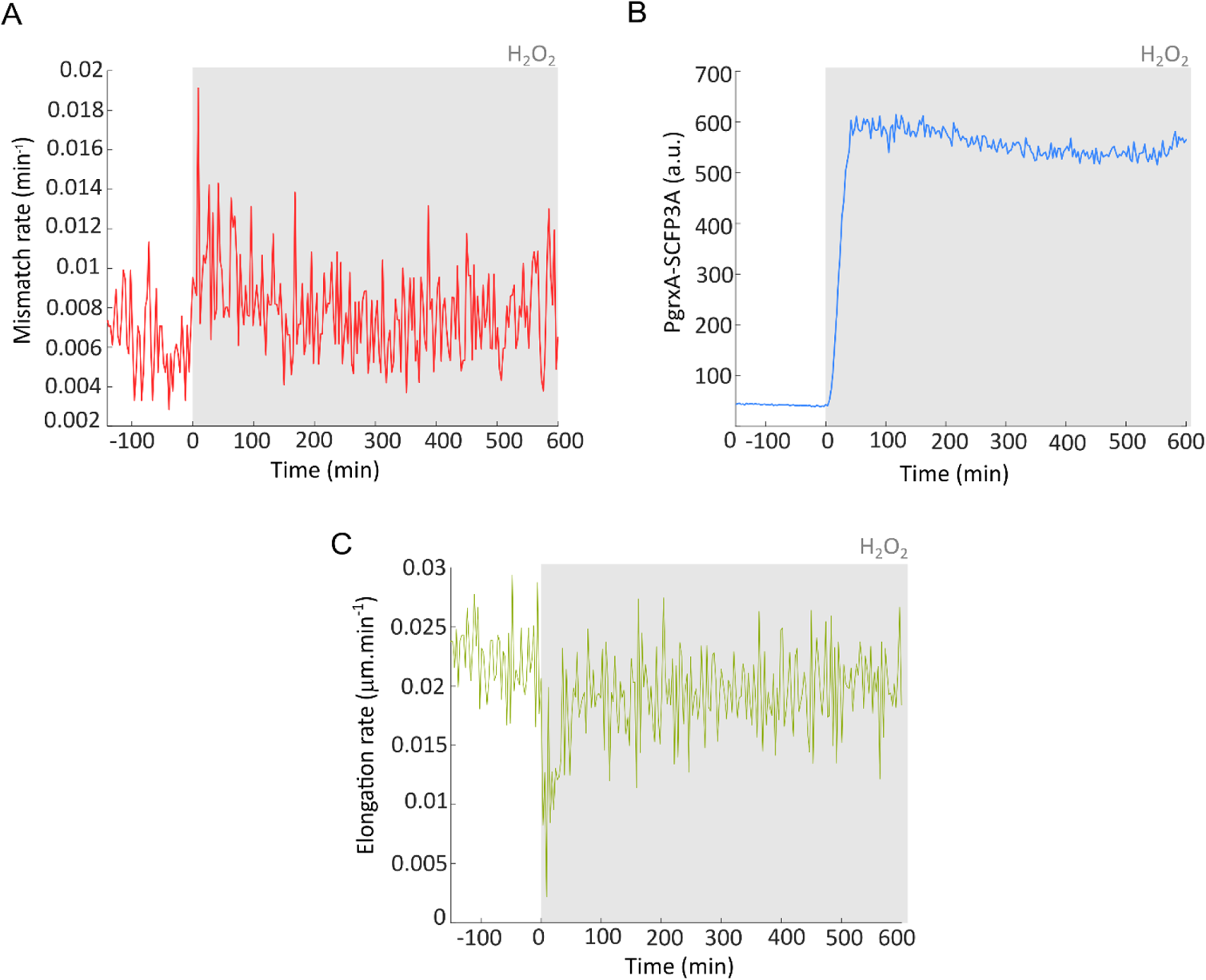
Similar mismatch rate and adaptation dynamics are observed in *E. coli* strain MG1655 compared to AB1157. (A) Rate of DNA mismatches per cell per minute before and during constant treatment with 100 µM H_2_O_2_(710 cells, 2 experiments) (B) PgrxA-SCFP3A expression before and during constant treatment with 100 µM H_2_O_2_ (710 cells, 2 experiments). (C) Cell elongation rate before and during constant treatment with 100 µM H_2_O_2_ (672cells, 2 experiments).

